# Experimental evolution reveals post-transcriptional regulation as a novel driver of *Leishmania* fitness gain

**DOI:** 10.1101/2021.03.22.436378

**Authors:** Laura Piel, K. Shanmugha Rajan, Giovanni Bussotti, Hugo Varet, Rachel Legendre, Caroline Proux, Thibaut Douché, Quentin Giai-Gianetto, Thibault Chaze, Thomas Cokelaer, Barbora Vojtkova, Nadav Gordon-Bar, Tirza Doniger, Smadar Cohen-Chalamish, Praveenkumar Rengaraj, Céline Besse, Anne Boland, Jovana Sadlova, Jean-François Deleuze, Mariette Matondo, Ron Unger, Petr Volf, Shulamit Michaeli, Pascale Pescher, Gerald F. Späth

**Affiliations:** Institut Pasteur, INSERM U1201, Unité de Parasitologie moléculaire et Signalisation, 75015 Paris, France; Université de Paris, 75013 Paris, France; The Mina and Everard Goodman Faculty of Life Sciences and Advanced Materials and Nanotechnology Institute, Bar-Ilan University, Ramat-Gan 52900, Israel; Institut Pasteur, Bioinformatics and Biostatistics Hub, Department of Computational Biology, USR 3756 IP CNRS, 75015 Paris, France; Institut Pasteur, Biomics, 75015 Paris, France; Institut Pasteur, UTechS MSBio, 75015 Paris, France; Institut Pasteur, Proteomics Platform Mass Spectrometry for Biology UTechS, C2RT, USR2000 CNRS, 75015 Paris, France; Department of Parasitology, Faculty of Science, Charles University, Prague 2, 128 44, Czech Republic; Université Paris-Saclay, CEA, Centre National de Recherche en Génomique Humaine, 91057, Evry, France

**Keywords:** *Leishmania*, experimental evolution, fitness, genomics, transcriptomics, post-transcriptional adaptation, small nucleolar RNAs

## Abstract

The protozoan parasite *Leishmania donovani* causes fatal human visceral leishmaniasis in absence of treatment. Genome instability has been recognized as a driver in *Leishmania* fitness gain in response to environmental change or chemotherapy. How genome instability generates beneficial phenotypes despite potential deleterious gene dosage effects is unknown. Here we address this important open question applying experimental evolution and integrative systems approaches on parasites adapting to *in vitro* culture. Phenotypic analyses of parasites from early and late stages of culture adaptation revealed an important fitness tradeoff, with selection for accelerated growth in promastigote culture (fitness gain) impairing infectivity (fitness costs). Comparative genomics, transcriptomics and proteomics analyses revealed a complex regulatory network driving parasite fitness, with genome instability causing highly reproducible, gene dosage-dependent changes in protein abundance linked to post-transcriptional regulation. These in turn were associated with a gene dosage-independent reduction in abundance of flagellar transcripts and a coordinated increase in abundance of coding and non-coding RNAs implicated in ribosomal biogenesis and protein translation. We correlated differential expression of small nucleolar RNAs (snoRNAs) with changes in rRNA modification, providing first evidence that *Leishmania* fitness gain in culture may be controlled by post-transcriptional and epitranscriptomic regulation. Our findings propose a novel model for *Leishmania* fitness gain in culture, where differential regulation of mRNA stability and the generation of fitness-adapted ribosomes may potentially filter deleterious from beneficial gene dosage effects and provide proteomic robustness to genetically heterogenous, adapting parasite populations. This model challenges the current, genome-centric approach to *Leishmania* epidemiology and identifies the *Leishmania* transcriptome and non-coding small RNome as potential novel sources for the discovery of biomarkers that may be associated with parasite phenotypic adaptation in clinical settings.

## Introduction

Parasitic protozoa of the genus *Leishmania* are the etiologic agents of a spectrum of severe diseases known as leishmaniases that cause substantial human morbidity and are among the five most serious parasitic diseases worldwide. Today, almost 1 billion people are at risk of *Leishmania* infection in close to 100 endemic countries throughout tropical and subtropical regions, with over 12 million people diagnosed with the infection [1]. Leishmaniasis represents a global public health challenge: recurrent epidemics are observed in South America, the Maghreb, Middle East, East Africa and India, and *Leishmania* infection has been declared an emerging disease in the EU and South East Asia [1, 2]. In absence of treatment, visceral leishmaniasis (VL - also known as Kala Azar) is the most severe and fatal form of the disease, caused either by *Leishmania* (*L.*) *donovani* or *L. infantum*.

Most *Leishmania* species show a digenetic life cycle comprising two major developmental stages that infect two distinct hosts. The motile, extracellular promastigote form of *Leishmania* proliferates inside the digestive tract of the sand fly insect vector. After migration towards the stomodeal valve, they eventually differentiate into the infectious metacyclic form, which is transmitted to the mammalian host during the blood meal. Once phagocytosed by host macrophages, metacyclic promastigotes differentiate in the non-motile, intracellular amastigote form that proliferates inside fully acidified, macrophage phagolysosomes of mammalian hosts. Aside stage differentiation, the success of *Leishmania* as a pathogenic microbe relies on its capacity to adapt to a variety of environmental fluctuations encountered in their hosts via an evolutionary process. Evolutionary adaptation relies on the classical Darwinian paradigm, where spontaneous mutations and stochastic changes in gene expression generate genetically and phenotypically heterogenous populations that compete for reproductive success in a given environment, thus driving natural selection of the fittest individuals [3]. While this process is well understood in viral and bacterial infections, only little information is available on evolutionary adaptation of eukaryotic pathogens, notably protozoan parasites. This is especially relevant to trypanosomatids, which - in contrast to classical eukaryotes - do not regulate expression of protein coding genes by transcriptional control. Transcription of protein coding genes in these early-branching eukaryotes is constitutive, with genes being arranged in long, polycistronic transcription units, and mature mRNAs being generated from precursors via a *trans*-splicing process unique to kinetoplastidae [4, 5]. In the absence of classical transcriptional regulation [5], *Leishmania* has evolved and emphasized other forms of gene expression control, notably regulation of RNA abundance by post-transcriptional regulation and gene dosage variations [6–10].

A hallmark of *Leishmania* biology is the intrinsic plasticity of its genome, with frequent copy number variations (CNVs) of individual genes or chromosomes linked to drug resistance or changes in tissue tropism [7, 11–16]. Combining experimental evolution and comparative genomics approaches, we recently linked both forms of genome instability to fitness gain *in vitro*. DNA read depth analysis of the genomes of *L. donovani* parasites adapting to culture identified amplification of a series of chromosomes as highly reproducible drivers of fitness gain [10]. Long-term adaptation in contrast was linked to the positive selection of gene copy number variants, which were amplified as part of functionally related, epistatic networks that allowed the emergence of phenotypes linked to ribosomal biogenesis, translation and proliferation [17]. *Leishmania* genomic adaptation thus occurs through a two-stage process reminiscent to other fast-growing eukaryotic cells (e.g. fungi and cancer cells [18, 19]), involving short-term adaptation by karyotypic changes and long-term adaptation through slower gene CNVs [10, 17, 20].

Together these reports draw a complex picture of *Leishmania* fitness gain in culture and raise a series of important new questions on (i) the nature of the genes that drive positive selection of the observed karyotypic changes during fitness gain *in vitro*, (ii) the potential fitness costs in infectivity associated with karyotypic adaptation, and (iii) the mechanisms evolved by the parasite to harness genome instability for fitness gain in culture and to compensate for deleterious gene dosage effects. Here we combined experimental evolution and integrative systems approaches to address these questions and gain novel insight into regulatory mechanisms underlying *Leishmania* fitness gain during adaptation to culture. Our analyses reveal mechanisms at gene, transcript and protein levels that harness genome instability for fitness gain *in vitro* through gene dosage-dependent changes that affect post-transcriptional regulation and gene dosage-independent changes in epitranscriptomic control and ribosomal biogenesis.

## Material and Methods

### Animals and ethics statement

Six to eight-week-old, female mice (*Mus musculus*, C57BL/6JRj) and 5 female Golden Syrian hamsters (*Mesocricetus auratus* RjHan:AURA, weighting between 60 – 70 g) were purchased from Janvier Laboratories. All animals were handled under specific, pathogen-free conditions in biohazard level 3 animal facilities (A3) accredited by the French Ministry of Agriculture for performing experiments on live rodents (agreement A75-15-01). Work on animals was performed in compliance with French and European regulations on care and protection of laboratory animals (EC Directive 2010/63, French Law 2013-118, February 6th, 2013). All animal experiments were approved by the Ethics Committee and the Animal welfare body of Institut Pasteur (dha190013 and 180091) and by the Ministère de l’Enseignement Supérieur, de la Recherche et de l’Innovation (project n°#19683).

### Parasites and culture

*Leishmania donovani* strain 1S2D (MHOM/SD/62/1S-CL2D) was obtained from Henry Murray, Weill Cornell Medical College, New York, USA and maintained by serial passages in hamsters. Amastigotes were recovered from infected hamster spleen and differentiated into promastigotes in M199 complete medium (M199, 10% FBS, 20 mM HEPES; 100 µM adenine, 2 mM L-glutamine, 10 µg/ml folic acid, 13.7 µM hemin, 4.2 mM NaHCO_3_, 1xRPMI1640 vitamins, 8 µM 6-biopterin, 100 units penicillin and 100 µg/ml streptomycin, pH 7.4) at 26°C. Promastigotes, derived from splenic amastigotes, were serially passaged once stationary phase was reached for less than 5 passages (EP, early passage) or 20 passages (LP, late passage) corresponding to ∼ 20 and 190 generations, respectively. Luciferase transgenic *Leishmania donovani* strain 1S (EP.luc, kindly provided by T. Lang; [21]) were serially passaged as described above.

### Experimental design

Strains issued from independent experimental evolution assays are identified by number (i.e. EP.1 and LP.1 are the strains resulting from experiment 1, see Figure S8 for details). For comparative analyses DNA, RNA or proteins were extracted from the different EP and LP logarithmic, stationary or metacyclic-enriched parasites as presented in Figure S8 and S9. Phenotypic characterization was performed on three different cultures prepared from EP.1, LP.1, EP.luc and LP.luc using frozen aliquots as starting material (see Figure S9).

### Parasite growth and determination of the generation time

Promastigotes in exponential growth phase were seeded at 2×10^6^ (EP) or 1×10^5^ (LP) parasites per ml in M199 complete medium. The different seeding densities allowed to compensate for the difference of EP and LP parasites in growth and to guarantee that they both reach stationary growth phase at the same time, which is essential for the comparative analyses of stationary-phase and metacyclic parasites. Parasites were counted every 24 hours and the generation time was calculated during logarithmic growth phase according to the formula *doubling time* = 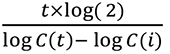. Experiments were performed in triplicates and statistical significance was assessed by t-test.

### Ficoll gradient centrifugation for metacyclic promastigote enrichment

EP and LP promastigote cultures were prepared as described above and maintained at stationary phase culture for 3 days when cell density, acidification and nutrition depletion trigger the differentiation from procyclic to metacyclic promastigotes. Parasites were collected and adjusted to 3×10^8^ cells/ml. Ficoll PM400 (GE Healthcare) was used to prepare a 20% stock solution in PBS and diluted for preparation of 10% and 5% Ficoll solutions. Four ml of 10% Ficoll were overlaid by 4 ml of 5% Ficoll and 4 ml of parasite suspension were layered on top of the Ficoll cushion. Tubes were centrifuged at 1,300 x *g* for 15 min at room temperature without brake. The metacyclic-enriched fractions were recovered at the interface between the 10% and 5% Ficoll layers. Parasites were washed with PBS and adjusted to the final concentration required for a given experiment.

### DNA extraction and sequencing

The different growth kinetics between EP and LP parasites were considered as described above and DNA was prepared from parasites in exponential culture phase. EP.1/LP.1, EP.7/LP.7, and LP.6 promastigotes were centrifuged at 1,600 x *g* for 10 min at room temperature. Approximately 1×10^8^ promastigotes from logarithmic growth phase were re-suspended in 200 µl PBS and genomic DNA was purified using DNeasy Blood and Tissue kit from Qiagen and RNase A according to the manufacturer’s instructions. DNA concentrations were measured in duplicate by fluorescence using a Molecular Device fluorescence plate reader (Quant-IT kits, Thermo Fischer Scientific). The quality of the DNAs was controlled determining the DNA Integrity Number (DIN) analyzing 20 ng of DNA on a TapeStation 4200 (Agilent). One µg genomic DNA was used to prepare a library for whole genome sequencing on an automated platform, using the Illumina “TruSeq DNA PCR-Free Library Preparation Kit”, according to the manufacturer’s instructions. After normalization and quality control, qualified libraries were sequenced on a HiSeqX5 platform from Illumina (Illumina Inc., CA, USA) at the Centre National de Recherche en Génétique Humaine (CEA, Evry, France), generating paired-ended, 150-bp reads. Sequence quality parameters were assessed throughout the sequencing run. Standard bioinformatics analysis of sequencing data was based on the Illumina pipeline to generate a FASTQ file for each sample.

### RNA extraction and sequencing

The different growth kinetics between EP and LP parasites were considered as described above. Total RNA was extracted from (i) EP.1, LP.1, EP.8, LP.8, EP.9 and LP.10 promastigotes at logarithmic culture phases (EP log/LP log), (ii) EP.1 and LP.1 parasites at 3 day-stationary culture phases (EP stat/LP stat), and (iii) metacyclic-enriched EP.1 parasites (EP.1 meta). Promastigotes were centrifuged at 3,000 x *g* for 10 min at room temperature and re-suspended in the lysis buffer supplied with the Qiagen RNeasy Plus kit. The samples were stored at -80°C and RNA extractions were performed according to the manufacturer’s instructions, including a DNase treatment. RNA integrity was controlled using the Agilent Bioanalyzer. DNase-treated RNA extracts were used for library preparation using the TruSeq Stranded mRNA sample preparation kit (Illumina, San Diego, California) according to the manufacturer’s instructions. An initial poly (A+) RNA isolation step (included in the Illumina protocol) was performed on total RNA to remove ribosomal RNA. Fragmentation was performed on the enriched fraction by divalent ions at high temperature. The fragmented RNA samples were randomly primed for reverse transcription followed by second-strand synthesis to create double-stranded cDNA fragments. No end repair step was necessary. An adenine was added to the 3’-end and specific Illumina adapters were ligated. Ligation products were submitted to PCR amplification. The obtained oriented libraries were controlled by Bioanalyzer DNA1000 Chips (Agilent, # 5067-1504) and quantified by spectrofluorimetry (Quant-iT™ High-Sensitivity DNA Assay Kit, #Q33120, Invitrogen). Sequencing was performed on the Illumina Hiseq2500 platform at the Biomics Center (Institut Pasteur, Paris, France) to generate single-ended, 130-bp reads bearing strand specificity.

For transcriptome-wide mapping of pseudouridine sites (Ψ-seq), total RNA from EP.1 and LP.1 parasites was either untreated or treated with N-cyclohexyl-N-β-(4-methylmorpholinium) (CMC) in bicine buffer (0.17 M CMC in 50 mM bicine, pH 8.3, 4 mM EDTA, 7 M urea) at 37°C for 20 min. Excess CMC was removed by ethanol precipitation. To remove CMC groups attached to G and U, the CMC-treated RNA was subjected to alkaline hydrolysis with Na_2_CO_3_ (50 mM, pH 10.4) at 37°C for 4h, as previously described [22–25]. The reacted RNA was recovered by phenol chloroform extraction, and ethanol precipitation. An adaptor was ligated to the 3’ end of the total RNA (upon fragmentation) before and after CMC treatment, and cDNA was prepared using AffinityScript reverse transcriptase (Agilent). The cDNA was then ligated to an adaptor, PCR amplified, and the samples were sequenced in an Illumina NextSeq machine in paired-end mode, 42-bp reads (20 million reads for each sample).

For the preparation of the small RNome, whole cell extracts were prepared from *L. donovani* EP and LP parasites (5×10^9^), that were washed with PBS and resuspended in 20 mM Tris-HCl (pH 7.7), 25 mM KCl, and 10 mM MgCl_2,_ were equilibrated in a nitrogen cavitation bomb (Parr Instruments Co.) with 750 psi N2 for 1h at 4°C, and disrupted by release from the bomb. After nitrogen cavitation, the ribonucleoproteins (RNPs) were extracted with 0.4 M KCl. Ribosomes were removed by centrifugation for 3h at 35,000 rpm in a Beckman 70.1Ti rotor (150,000 x *g*) and the supernatant was defined as post-ribosomal supernatant (PRS). RNA was extracted after treatment with 100 µg/ml of Proteinase K, 1% SDS in the presence of 100 µg/ml DNaseI and was used for library preparation as described previously [23]. The samples were sequenced in an Illumina NextSeq machine in paired-end mode, 42-bp reads (40 million reads for each sample).

### Protein extraction, digestion and LC-MS/MS acquisition

Exponentially growing EP.2, EP.3, EP.4, EP.5 and LP.2, LP.3, LP.4 and LP.5 promastigotes were centrifuged at 1600 x *g* for 10 min at 4°C and washed three times with cold PBS. Parasite lysates were prepared in 8 M urea, 50 mM Tris, supplemented with a protease inhibitor cocktail (cOmplete™ from Roche) and a phosphatase inhibitor cocktail (PhosStop from Roche), 1 ml of lysis buffer per 1.5×10^9^ promastigotes. After 10 min incubation at 4°C followed by sonication for 5 min (sequence of 10s pulse and 20s pause) the lysates were centrifuged 15 min at 14,000 x *g*, 4°C and the supernatant was collected and stored at -80°C until use. Proteins were quantified by RC DC™ protein assay (Bio-Rad) and a control of the protein pattern of all the extracts was performed by SDS-PAGE and silver staining. All the biological samples were further processed for MS-based proteomics approach, data acquisition, and statistical analyses.

Biological samples were adjusted to 1.3 µg.µl^-1^ in lysis buffer. Disulfide bridges were reduced in 5 mM DTT (Sigma - 43815) for 30 min and alkylated in 20 mM iodoacetamide (Sigma - I1149) for 30 min at room temperature in the dark. Protein samples were diluted 10-fold in 50 mM Tris-HCl and digested with Sequencing Grade Modified Trypsin (Promega - V5111) at a Protein:Trypsin ratio 50:1 overnight. Then a second digestion was performed to complete this step. Proteolysis was stopped by adding formic acid (FA, Fluka - 94318) at a 1% final concentration. Resulting peptides were desalted using Sep-Pak SPE cartridge (Waters) according to the manufacturer’s instructions. Peptides were concentrated to almost dryness and were resuspended in 2% Acetonitrile (ACN) / 0.1% FA just before LC-MS/MS injection.

All analyses were performed on a Q Exactive™ Plus Mass Spectrometer (Thermo Fisher Scientific) coupled with a Proxeon EASY-nLC 1000 (Thermo Fisher Scientific). One µg of peptides was injected into a home-made 50 cm C18 column (1.9 μm particles, 100 Å pore size, ReproSil-Pur Basic C18, Dr. Maisch GmbH, Ammerbuch-Entringen, Germany). Column equilibration and peptide loading were performed at 900 bars in buffer A (0.1% FA). Peptides were separated with a multi-step gradient of 2 to 5% buffer B (80% ACN, 0.1% FA) for 5 min, 5 to 22% buffer B for 150 min, 22 to 45% buffer B for 60 min, 45 to 80% buffer B for 10 min at a flow rate of 250 nL/min over 240 min. Column temperature was set to 60°C. MS data were acquired using Xcalibur software using a data-dependent method. MS scans were acquired at a resolution of 70,000 and MS/MS scans (fixed first mass 100 m/z) at a resolution of 17,500. The AGC target and maximum injection time for the survey scans and the MS/MS scans were set to 3E^6^, 20 ms and 1E^6^, 60ms respectively. An automatic selection of the 10 most intense precursor ions was activated (Top 10) with a 45s dynamic exclusion. The isolation window was set to 1.6 m/z and normalized collision energy fixed to 28 for HCD fragmentation. We used an underfill ratio of 1.0% corresponding to an intensity threshold of 1.7E^5^. Unassigned precursor ion charge states as well as 1, 7, 8 and >8 charged states were rejected and peptide match was disabled.

### Data analyses

#### WGS analysis

Genomic DNA reads were aligned to the *L. donovani* Ld1S reference genome (https://www.ncbi.nlm.nih.gov/bioproject/PRJNA396645<u>, GCA_002243465.1</u>) with BWA mem (version 0.7.12) with the flag -M to mark shorter split hits as secondary. Samtools fixmate, sort, and index (version 1.3) were used to process the alignment files and turn them into bam format [26]. RealignerTargetCreator and IndelRealigner from the GATK suite were run to homogenize indels [27]. Eventually, PCR and optical duplicates were labeled with Picard MarkDuplicates [version 1.94(1484)] (https://broadinstitute.github.io/picard/) using the option “VALIDATION_STRINGENCY=LENIENT”. For each read alignment file, Samtools view (version 1.3) and BEDTools genomecov (version 2.25.0) were used to measure the sequencing depth of each nucleotide [28]. Samtools was run with options “-q 50 -F 1028” to discard reads with a low map quality score or potential duplicates, while BEDTools genomecov was run with options “-d -split” to compute the coverage of each nucleotide. The coverage of each nucleotide was divided by the median genomic coverage. This normalization is done to account for library size differences. The chromosome sequencing coverage was used to evaluate aneuploidy between EP.1 and LP.1 samples. Then for each sample and for each chromosome, the median sequencing coverage was computed for contiguous windows of 2,500 bases. As previously published [10], the stably disomic chromosome 36 was used to normalize chromosome read depth and to estimate chromosome polysomy levels in each sample. Gene counts were produced using featureCounts (version 1.4.6-p3 [29]) with these parameters: -s 0 -t gene -g gene_id and were normalized according to the median-ratio method.

#### Genome binning

The reference genome was divided into contiguous windows of a fixed length, and the sequencing coverage of each window was evaluated and compared across different samples. A window length of 300 bases was used for the shown scatter plot assessing genome-wide CNVs. Both the mean sequencing coverage normalized by the median chromosome coverage and the mean read MAPQ value were computed [20].

#### RNAseq analysis

For total RNAseq data, the bioinformatics analysis was performed using the RNA-seq pipeline from Sequana [30]. Reads were cleaned of adapter sequences and low-quality sequences using cutadapt version 1.11 [31]. Only sequences of at least 25 nucleotides in length were considered for further analysis. STAR version 2.5.0a [32], with default parameters, was used for alignment on the reference genome (GCA_002243465.1). Genes were counted using featureCounts version 1.4.6-p3 [29] from Subreads package (parameters: -t gene -g gene_id -s 1). Count data were analyzed using R version 3.6.1 [33] and the Bioconductor package DESeq2 version 1.24.0 [34]. The normalization and dispersion estimation were performed with DESeq2 using the default parameters and statistical tests for differential expression were performed applying the independent filtering algorithm. For each pairwise comparison, raw p-values were adjusted for multiple testing according to the Benjamini and Hochberg (BH) procedure [35] and genes with an adjusted p-value lower than 0.01 were considered differentially expressed. The RNAseq data have been deposited in NCBI’s Gene Expression Omnibus [36] and are accessible through GEO Series accession number GSE165615 (https://www.ncbi.nlm.nih.gov/geo/query/acc.cgi?acc=GSE165615).

For small RNome analysis, Ψ-seq and detection of pseudouridylated sites, the 42 bp sequence reads obtained from the Illumina Genome Analyzer were first trimmed of Illumina adapters using the FASTX toolkit (http://hannonlab.cshl.edu/fastx_toolkit), and reads of 15 nucleotides or less were discarded from subsequent analysis. The remaining reads were mapped to the reference genome (GCA_002243465.1) using SMALT v0.7.5 (https://www.sanger.ac.uk/tool/smalt-0/) with the default parameters. Only properly paired partners were retained. Each read pair was “virtually” extended to cover the area from the beginning of the first read to the end of its partner. For each base, the number of reads initializing at that location as well as the number of reads covering the position were calculated. A combination of BEDTools v2.26.0 Suite (http://bedtools.readthedocs.io/en/latest/) and in-house Perl scripts was used to calculate the Ψ-ratio and Ψ-fc (fold change), as previously described [23, 24].

#### Proteomics analysis

Raw data were analyzed using MaxQuant software version 1.5.3.8 [37] using the Andromeda search engine [38]. The MS/MS spectra were searched against the Ld1S database (https://www.ncbi.nlm.nih.gov/bioproject/PRJNA396645<u>, GCA_002243465.1</u>). The settings for the search included (i) trypsin digestion with a maximum of two missed cleavages, (ii) variable modifications for methionine oxidation and N-terminal acetylation, and (iii) fixed modification for cysteine carbamidomethylation. The minimum peptide length was set to 7 amino acids and the false discovery rate (FDR) for peptide and protein identification was set to 0.01. The main search peptide tolerance was set to 4.5 ppm and to 20 ppm for the MS/MS match tolerance. The setting ‘second peptides’ was enabled to identify co-fragmentation events. Quantification was performed using the XIC-based Label-free quantification (LFQ) algorithm with the Fast LFQ mode as previously described [39]. Unique and razor peptides, including modified peptides, with at least two ratio counts were accepted for quantification. The mass spectrometry proteomics data were deposited to the ProteomeXchange Consortium via the PRIDE partner repository with the dataset identifier PXD020236 [40].

For the differential analyses, proteins categorized as ‘reverse’, ‘contaminant’ and ‘only identified by site’ were discarded from the list of identified proteins. After log2 transformation, LFQ values were normalized by median centering within conditions (*normalizeD* function of the R package *DAPAR* [41]). Remaining proteins without any LFQ value in one of the conditions (either EP or LP) and at least two values in the other condition were considered as exclusively expressed proteins. Missing values across the four biological replicates were imputed using the imp.norm function of the R package norm (norm: Analysis of multivariate normal datasets with missing values. 2013 R package version 1.0-9.5). A limma t test was applied to determine proteins with a significant difference in abundance while imposing a minimal fold change of 2 between the conditions to conclude that they are differentially abundant [42, 43]. An adaptive Benjamini-Hochberg procedure was applied on the resulting p-values using the function *adjust.p* of R package *cp4p* [44] and the robust method described in Pounds et al. [45] to estimate the proportion of true null hypotheses among the set of statistical tests. The proteins associated to an adjusted p-value inferior to a False Discovery Rate (FDR) of 0.01 have been considered as significant and differentially abundant proteins.

#### Gene Ontology (GO)-enrichment analyses and gene category assignment

The Biological Networks Gene Ontology tool (BiNGO) plugin of the Cytoscape software package (version 3.8.2) was used. A Benjamini & Hochberg false discovery rate with a significance level of 0.05 was applied. The lists of *L. donovani* GO terms were built in house (see Script in Supplementary data). In order to assign the Gene Ontology Identifiers (GO IDs) we combined the GO-derived identifiers with the ones available from the corresponding orthologs in target species: LdBPK, *L. infantum*, *L. major*, *L. mexicana*, *Typanosoma brucei brucei 927* (Tbru) and *Typanosoma cruzi* (Tcru). For each target species we retrieved both the “curated” and “computed” GO IDs from TriTrypDB on the 11/09/2019. OrthoFinder with the DIAMOND search program was applied to establish orthology between the genes in Ld1S and in target species. In “one-to-many” orthology relations we concatenated all the non-redundant GO IDs from all the homologs. The GO IDs were then assigned based on the hierarchy: LdBPK curated > LdBPK GO > *L. infantum* curated > *L. major* curated > *L. mexicana* curated > Tbru curated > Tcru curated > LdBPK computed > *L. infantum* computed > *L. major* computed > *L. mexicana* computed > Tbru computed > Tcru computed. The GO IDs were assigned if not present in any higher rank GO ID data set. The GO IDs of snoRNAs, UsnRNA, SLRNA and 7SL classes defined by homology with *L. major* Friedlin genes were manually attributed. Overall, we assigned biological process (BP), molecular function (MF) and cellular component (CC) GO IDs to 5,246, 4,521 and 7,236 Ld1S genes [17] (Supplementary data S1).

Total frequency represents the percentage of genes associated with a given GO term in the genome compared to the total number of annotated genes. Cluster efficiency represents the percentage of genes for a given GO term in a data set compared to all genes that are annotated with any GO term in the same data set. Enrichment score corresponds to the number of genes for a given GO term in a data set compared to the total number of genes sharing the same GO identifier in the genome. Cluster efficiencies, total frequencies and enrichment scores are shown in tables 2 to 6 in the GO analyses sections.

Genes and proteins were assigned to categories by combining GO analysis and manual inspection for annotations. Genes or proteins annotated for a GO term, a known function or product were considered to estimate the percentage of genes in each category. Gene or protein categories are presented in tables 2 to 6.

### Northern blot analyses

Total RNA extracted from EP and LP cells (10 µg) were separated on 10% acrylamide denaturing gels, transferred to nitrocellulose membranes and analyzed by autoradiography. RNA probes were prepared by *in vitro* transcription using α-^32^P-UTP [23]. Three independent northern blots were performed.

### Bone marrow-derived macrophages and infection

Bone marrow exudate cells were recovered from tibias and femurs of C57BL/6JRj female mice (Janvier Labs) and macrophages differentiated in DMEM complete medium (DMEM, 15% FBS, 10 mM HEPES, 50 µM 2-mercaptoethanol, 50 units of penicillin and 50 µg/ml of streptomycin) supplemented with 75 ng/ml of recombinant mouse colony stimulating factor-1 (rmCSF-1, ImmunoTools) [46]. A total of 1.5×10^5^ bone marrow-derived macrophages (BMDMs) were plated on glass coverslips in 24-wells plates and incubated overnight at 37°C, 5% CO_2_ prior to *Leishmania* infection.

Promastigotes from stationary culture phase or metacyclic-enriched parasite fractions were pelleted by centrifugation at 3,000 x *g* for 10 min at room temperature and re-suspended in PBS. The concentration was adjusted to 6×10^7^ parasites per ml and 50 µl were added to the BMDM cultures at a multiplicity of infection (MOI) of 20 parasites per 1 macrophage. Plates were centrifuged at 300 x *g* for 5 min at room temperature to allow for a faster sedimentation of the parasites onto the macrophage monolayer. After 2h of contact, coverslips were washed by successive baths in pre-warmed PBS to remove extracellular parasites and transferred into new 24-wells plates containing fresh pre-warmed DMEM culture medium supplemented with 30 ng/µl of rmCSF-1. At 4, 24, 48 and 168h post-infection, cells were fixed in 4% paraformaldehyde (Electron Microscopy Science) and macrophage and parasite nuclei were stained with Hoechst 33342. Images were acquired using a Zeiss Apotome microscope at 40x magnification connected to an Axiocam camera. All the infections were performed in triplicates and at least 100 macrophages were counted per coverslip. The total numbers of infected and non-infected macrophages were recorded and the percentage of infection, the number of parasites per 100 cells and the number of parasites per infected macrophages was calculated and normalized to the values obtained at the initial 4-hour time point. The replication rate in macrophages was calculated between day 1 and day 6 after infection. All the experiments were performed three times in triplicates (see experimental overview in Figure S9 for details) using independent preparations of primary macrophages for each infection.

### Morphological analyses

Parasites were seeded on poly L-lysine treated coverslips and fixed in 2.5% glutaraldehyde. Coverslips were mounted on glass slides using Mowiol® 4-88 (Sigma-Aldrich). Images were acquired using an Axiophot microscope at 63x magnification and an Andor camera. Length and width of the parasite cell body, and flagellum length were measured for at least 200 promastigotes using the Image J Fidji software package (https://imagej.net/). The ratios flagellum over body length and body length over body width were determined for the 200 parasites and the Kruskal-Wallis test was used for statistical analysis. The experiment was performed in duplicate.

### Sand fly infection

The colony of *Phlebotomus orientalis* (originating from Ethiopia), the natural vector of *L. donovani*, was maintained in the insectary of the Department of Parasitology, Charles University in Prague, under standard conditions (26°C on 50% sucrose and 14h light/10h dark photoperiod) as described previously [47, 48].

Promastigotes from logarithmic-phase cultures (day 3-4 in culture) were washed twice in saline solution and resuspended in heat-inactivated rabbit blood at a concentration of 1×10^6^ promastigotes/ml. Sand fly females (5-9 days old) were infected by feeding through a chick-skin membrane (BIOPHARM) on a promastigote-containing suspension. Engorged sand flies were separated and maintained under the same conditions as the colony. On day 8 post-blood meal (PBM), 150 sand fly females were dissected. The thoracic parts and abdominal parts of infected guts were collected separately and pooled together into two samples: thoracic parts of gut (TP) and abdominal parts of gut (AP). The exact numbers of all parasite stages were calculated using a Burker apparatus and the proportion of metacyclic forms was identified on a Giemsa-stained smears separately for TP and AP. *Leishmania* with flagellar length < 2 times body length were scored as procyclic forms and those with flagellar length ≥2 times body length as metacyclic forms [49].

## Results

### *L. donovani* long-term culture adaptation causes a fitness tradeoff between in vitro proliferation and infectivity

In microbial culture, fitness gain (defined as reproductive capacity) largely equals the level of cell proliferation. Adaptation to *in vitro* growth thus represents a simple experimental system to assess mechanisms underlying fitness gain. Here we applied such an experimental evolution approach on *L. donovani* amastigotes isolated from infected hamster spleen. Derived promastigotes at early-passage (EP.1) and late-passage (LP.1) were monitored for growth and infectivity with the aim to assess regulatory mechanisms underlying fitness gain and fitness cost observed during culture adaptation. Analyzing cell growth during promastigote culture adaptation revealed robust fitness gain as judged by the reduction in generation time from 13.76 +/- 1.18 hours for EP.1 to 9.76 +/- 0.93 hours for LP.1 promastigotes (Figure 1A). We next evaluated fitness of these parasites in intracellular macrophage infection, where reproductive success depends on parasite resistance to host cell cytolytic activities, amastigote differentiation and proliferation. BMDMs were incubated with EP.1 and LP.1 promastigotes from day-3 stationary culture (referred to as EP.1 stat and LP.1 stat) and intracellular growth was monitored microscopically for 7 days as previously described [50]. Even though the number of EP.1 stat and LP.1 stat intracellular parasites decreased over the first 24h post-infection, only EP.1 parasites recovered and established persistent infection, while the number of LP.1 parasites steadily declined during the subsequent 6 days (Figure 1B and Figure S1A). The same results were obtained in an independent evolutionary experiment conducted with transgenic parasites expressing luciferase, EP.luc and LP.luc (Figure S1C). Together these data firmly establish the highly reproducible nature of the fitness tradeoff between *in vitro* proliferation and infectivity in macrophages as a result from long-term *L. donovani* culture adaptation and confirm our previous reports [10, 50].

**Figure 1:**
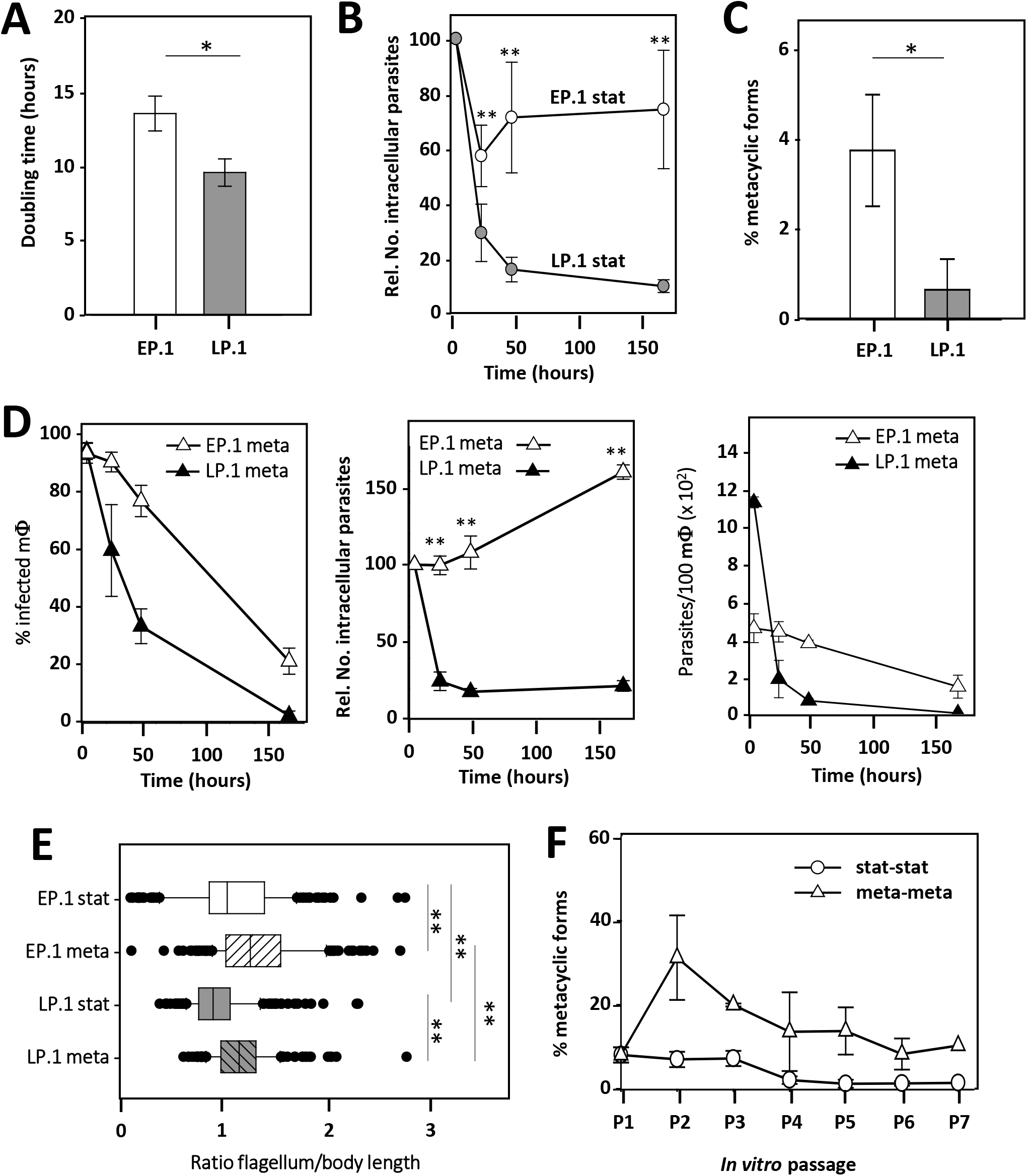
Phenotypic analysis of EP.1 and LP.1 parasites reveals fitness tradeoff between in vitro proliferation and macrophage infectivity. (A) Histogram plot representing the generation time of EP.1 and LP.1 promastigotes in culture calculated based on parasite density during logarithmic growth phase. The mean value of three independent experiments +/- SD is represented. *p-value ≤ 0.05. (B) Macrophage infection assay. The mean relative number of intracellular EP.1 (open circles) and LP.1 (grey circles) parasites +/- SD of three independent triplicate experiments using promastigotes from day-3 stationary culture is represented. **p-value ≤ 0.01. (C) Histogram plot representing the percentage of EP.1 and LP.1 metacyclic forms that were enriched by Ficoll density gradient centrifugation from cultures at stationary growth phase. Each bar represents the mean +/- SD of four independent experiments. *p-value ≤ 0.05. (D) Macrophage infection assay using Ficoll-enriched metacyclic parasites. Percentage of infected macrophages (left panel), mean relative number of intracellular EP.1 and LP.1 parasites (middle panel) and mean number of parasites per 100 macrophages (right panel) are shown. Open triangles, EP.1 meta; close triangles, LP.1 meta. The mean values +/- SD of one triplicate experiment are shown. **p-value ≤ 0.01. (E) Morphological characterization of EP.1 and LP.1 Ficoll-enriched metacyclic parasites. Body width, flagellum and body length were measured on 200 promastigotes using the Image J software package. The ratio flagellum-to-body length was computed from two biological replicate experiments and the median values +/- SD are represented by the box plot with the upper and lower quartiles indicated. **p-value ≤ 0.01. (F) Percentage of metacyclic-like parasites recovered by Ficoll gradient centrifugation from cultures seeded successively for 6 *in vitro* passages with either EP.1 from stationary growth phase (stat-stat) or EP.1 metacyclic-enriched parasites (meta-meta). Mean values of two independent experiments are shown with +/-SD denoted by the bars.

We then tested if the fitness cost of LP.1 stat in infectivity was due to a differentiation defect of infectious metacyclic promastigotes. Considering that stationary phase cultures are composed of different forms of promastigotes, a Ficoll gradient centrifugation method was used to enrich and quantify metacyclic parasites. This method, based on separation of the different parasite forms according to their density [51], allowed to reveal a 5.5-fold reduction in the number of metacyclic parasites from 3.8% in EP.1 stat to 0.69% in LP.1 stat cultures (Figure 1C), the latter one in addition being compromised to establish macrophage infection (Figure 1D and S1B). These results document that the fitness cost in LP.1 meta not only affects the quantity but also the quality of differentiating metacyclic parasites. This was further confirmed by their atypical morphology that was different to *bona fide*, sand fly-isolated metacyclic promastigotes (Figure S2A), corresponding to leptomonad-like forms as judged by flagellum/body-length ratio and body shape [52, 53] (Figure 1E, Figure S2A-C). Surprisingly, unlike observed when passaging EP.1 stat parasites, metacyclogenesis was maintained in cultures that were passaged using metacyclic-enriched parasites (EP.1 meta) (Figure 1F).

### Transcriptome profiling informs on mechanisms underlying fitness tradeoff

We performed RNA-seq analyses using poly (A+)-enriched mRNA obtained from three replicates of EP.1 and LP.1 log, stat, and EP.1 meta parasites. The low yield in LP.1 meta parasites precluded their analysis by RNA-seq. Principal component and hierarchical clustering analyses demonstrated that transcript profiles of EP.1 and LP.1 parasites grouped according to stage, indicating that stage-specific expression changes in log, stat and meta parasites dominate over those associated with the EP.1/LP.1 promastigote fitness tradeoff (Supplementary Table 2-a to -f, Figure 2A, Figure S3A). Significant stage-specific changes were observed in EP.1 and LP.1 parasites during the log-stat transition for respectively 54.2% and 49.3% of the transcripts and ca. 35% of the promastigote transcriptome was modulated between EP.1 stat and EP.1 meta (Figure 2B and S3B, Supplementary Table 2-a to -f). As expected from the increased motility described for metacyclic parasites, we indeed observed increased abundance in EP.1 meta compared to EP.1 log and EP.1 stat for respectively 48 and 51 genes linked to motility and flagellar biogenesis (Supplementary Table 2-p).

**Figure 2:**
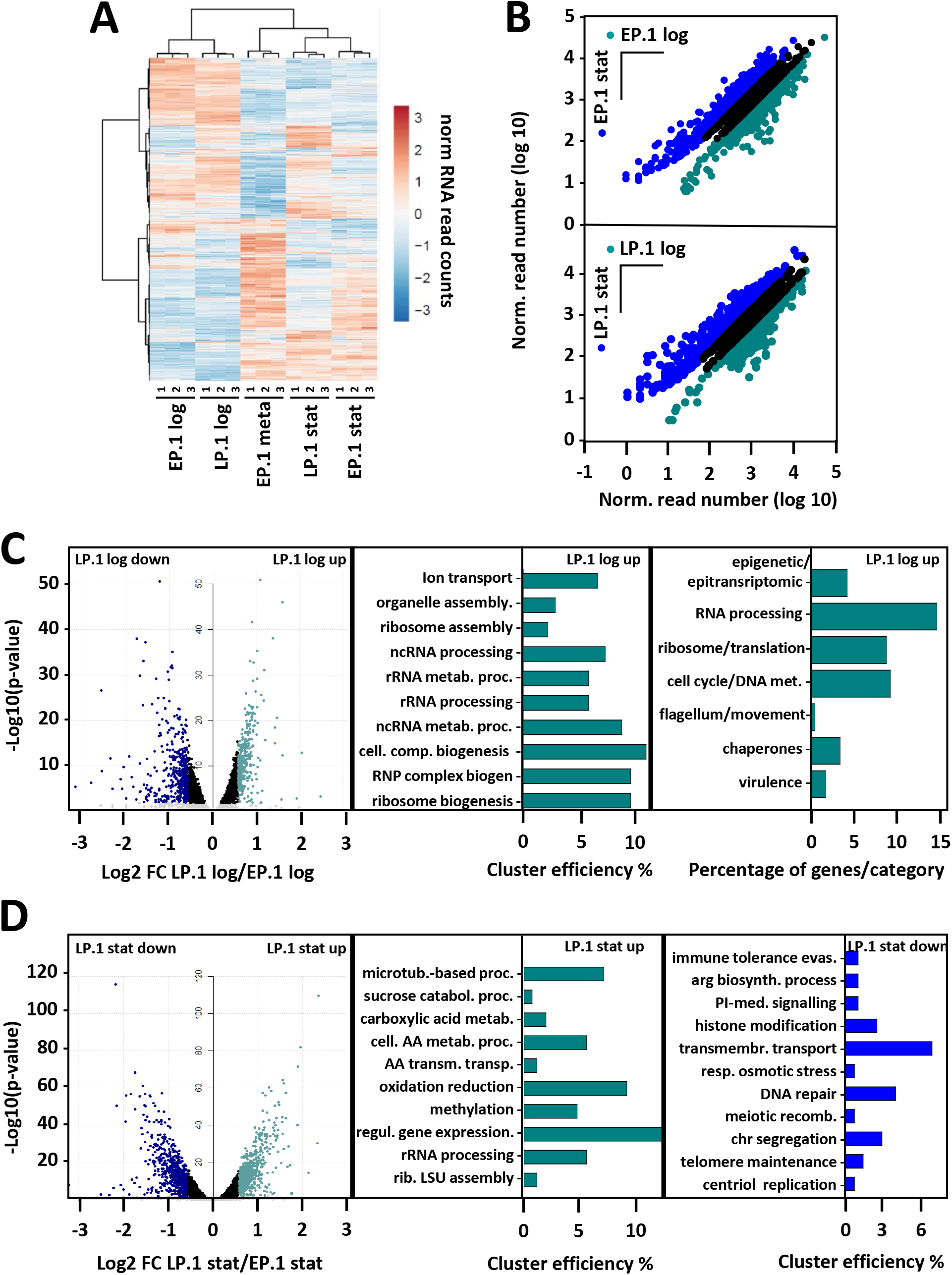
RNA-seq analyses of EP.1 and LP.1 promastigotes reveal stage-specific changes in RNA abundance and RNA signatures linked to fitness gain in culture and fitness cost in infectivity. (A) Cluster analysis of differentially expressed genes observed in triplicate RNAseq analyses of EP.1 log and LP.1 log, EP.1 stat and LP.1 stat, and EP.1 meta parasites. (B) Ratio plots of normalized RNAseq reads for EP.1 log compared to EP.1 stat (upper panel) and LP.1 log compared to LP.1 stat (lower panel). Blue and dark cyan dots represent gene expression changes with FC > 1.5 and adjusted p-value ≤ 0.01; black dots correspond to gene expression changes with adjusted p-value > 0.01. Only genes with at least 10 reads in one of the two conditions were considered. Top panel, 1499 and 1501 transcripts more abundant in EP.1 log (dark cyan) and EP.1 stat (blue), respectively. Lower panel, 1129 and 1384 transcripts more abundant in LP.1 log (dark cyan) and LP.1 stat (blue), respectively (see Supplementary table 2-a and -d). (C) Differential expression profiling of LP.1 log and EP.1 log parasites. Transcripts more abundant in EP.1 log correspond to transcripts less abundant in LP.1 log. Volcano plot representing the changes in transcript abundances of LP.1 log and EP.1 log parasites with 344 transcripts more abundant in LP.1 log (LP.1 log up) versus 433 transcripts less abundant in LP.1 log (LP.1 log down) (left panel) (see Supplementary table 2-g and -h for the list of regulated genes). Transcripts with significant increased abundance FC > 1.5 and adjusted p-value ≤ 0.01 in LP.1 log up and LP.1 log down are indicated respectively in cyan and blue and were used to perform the GO analysis for the category ‘biological process’. The histogram plot (middle panel) shows ‘cluster efficiency’, which represent the percentage of genes associated with a given GO term compared to the total number of genes with any GO annotation in the considered set of genes. Only functional enrichments associated with adj. p-value < 0.05 were considered. For transcripts more abundant in LP.1 log (LP.1 log up), only 134 out of 344 genes are associated with a GO ID (see Supplementary table 2-o for details). Transcripts showing increased abundance and adj. p-value <0.01 in LP.1 log were categorized in functional groups (right panel). The histogram plot shows the percentage of genes which represent the number of genes for the indicated gene families compared to the total number of genes with a known function or product (see Supplementary table 2-i for details). (D) Differential expression profiling of LP.1 stat and EP.1 stat parasites. Transcripts more abundant in EP.1 stat correspond to transcripts less abundant in LP.1 stat. Volcano plot representing the changes in transcript abundances of LP.1 stat and EP.1 stat parasites with 662 transcripts more abundant in LP.1 stat (LP.1 stat up) versus 710 transcripts less abundant in LP.1 stat (LP.1 stat down) (left panel) (see Supplementary table 2-k and -l for the list of up regulated genes). Transcripts with significant increased abundance FC > 1.5 and adjusted p-value ≤ 0.01 in LP.1 stat up and LP.1 stat down are indicated respectively in cyan and blue and were used to perform the GO analysis. Results of GO analyses for the category ‘biological process’ performed on transcripts showing statistically significant increased (middle panel) and decreased (right panel) abundance in LP.1 stat are shown (see Supplementary table 2-o). Cluster efficiencies were calculated based on 258 and 274 genes with GO IDs in LP.1 stat up and LP.1 stat down set of genes, respectively. Only the functional enrichments associated with adj. p-value < 0.05 were considered.

We next assessed changes in transcript abundance observed at logarithmic growth phase in LP.1 compared to EP.1 promastigotes to gain first insight into pathways associated with *in vitro* fitness gain (i.e. accelerated growth). We identified 344 transcripts with significantly increased abundance in LP.1 log (Figure 2C, left panel, Supplementary Table 2-g) and revealed functional enrichment in this dataset for the GO terms ‘ribosome biogenesis’, ‘ribosome assembly’, and ‘rRNA processing’ (Figure 2C, middle panel, Figure S3E and Supplementary Table 2-o). Combining GO analysis and manual inspection of gene annotation, 56 genes fell in the categories RNA processing and ribosome/translation, representing 24% of the quantified genes that are annotated for a known function or product (Figure 2C, right panel, Supplementary Table 2-i). LP.1 log fitness gain in culture thus likely reflects an increase in translation efficiency, which may allow for accelerated growth observed in these cells. Further analysis revealed increased abundance of other transcripts implicated in various regulatory processes linked to proliferation (Figure 2C, right panel, Supplementary Table 2-i), including epigenetic/epitranscriptomic regulation (10 genes, e.g. Ld1S_110036500 encoding for a Pseudouridylate synthase 10, Ld1S_260334600 encoding for a RNA pseudouridylate synthase and Ld1S_330597500 encoding for a Histone methyltransferase DOT1) and cell cycle/DNA metabolism (22 genes, e.g. Ld1S_050817000 encoding for CYC2-like cyclin, or Ld1S_330603400 encoding for the cell division control protein CDC45) (see Supplementary Table 2-i for more examples). To assess the reproducibility of these results, we performed RNA-seq analysis on independently evolved LP and EP log parasites (EP.8, EP.9 and LP.8, LP.10) (see Figure S8 for details). Just like in the EP.1/LP.1 comparison, enrichment was observed for various categories linked to ribosomal biology, thus confirming the link between *in vitro* fitness gain and protein translation.

In contrast, no GO enrichment was observed for the 433 transcripts showing significant reduced abundance in LP.1 log (Supplementary Table 2-h). Manual inspection of gene annotations identified various pathways implicated in metabolism and energy production (e.g. genes encoding for respiratory chain proteins, amino acid and sugar metabolism, fatty acid biosynthesis), signaling (numerous kinases and phosphatases) and flagellum/motility (including four genes encoding for paraflagellar rod components) (Figure S3C, Supplementary Table 2-j). An even stronger reduction of transcripts associated with motility was found in our second transcriptomic analysis of independently ovolved LP and EP log parasites (EP.8, EP.9 and LP.8, LP.10, see Figure S8 for details) (Figure S3D, Supplementary Table 2bis -b and -c). These pathways suggest a potential retooling of the LP.1 log energy metabolism in the nutrition-rich culture environment, and selection against motility, which is not essential in culture and may liberate the energy required for faster growth. Surprisingly, one of the most significant decreases in transcript abundance in these cells was observed for a gene encoding for a 5S ribosomal RNA, along four other genes encoding for ribosomal components (Supplementary Table 2-h), even though other ribosomal components were upregulated in LP.1 log. This result provided a first indication that LP.1 log fitness gain in culture not only depends on the quantity, but likely also the quality or type of ribosomes, e.g. their ribonucleoprotein composition, which may control the fitness-adapted expression profile at the translational level, for example by changing the ribosome translation specificity or efficiency.

Finally, we assessed changes in transcript abundance observed at stationary growth phase in LP.1 compared to EP.1 promastigotes to gain further insight into mechanisms of fitness loss (i.e. attenuated infectivity). We identified 662 transcripts with significantly increased abundance in LP.1 stat (Figure 2D, left panel, Supplementary Table 2-k). Enrichment was observed for the GO terms ‘ribosomal large subunit assembly’, ‘rRNA processing’ and ‘regulation of gene expression’ (Figure 2D, middle panel, Figure S3F upper panel, Supplementary Table 2-o). In contrast, LP.1 stat promastigote showed reduced abundance for 710 transcripts, including transcripts linked to the GO terms ‘histone modification’, ‘DNA repair’, ‘transmembrane transport’ (Figure 2D, right panel, Supplementary Table 2-o) and fifteen transcripts manually associated to cell cycle (Supplementary Table 2-n). Likewise, decreased abundance was observed for transcripts associated with the GO term ‘evasion or tolerance of immune response of other organism involved in symbiotic interaction’ and ‘virulence’. Manual inspection allowed us to enrich this last term from originally three to 29 genes (Figure 2D, right panel and Figure S3G and Supplementary Table 2-o and -n). Indeed, almost 14% of the transcripts with reduced abundance correspond to genes previously associated with parasites infectivity, including GP63 as well as 31 amastin surface glycoproteins and amastin-like proteins (Figure S3G and Supplementary Table 2-n). Hence, the reduced expression in LP.1 stat parasites of these genes could be associated with attenuated infectivity we observed in LP.1 stat and meta parasites (see Figure 1B and D) [54, 55].

In conclusion, our data link increased fitness in *in vitro* growth of LP.1 log to a gain-of-function phenotype associated with proliferation, ribosomal biogenesis, and translation. Conversely, the reduced fitness in infectivity of LP.1 stat was associated with a loss-of-function phenotype linked to decreased expression of virulence genes.

### Post-transcriptional adaptation during promastigote fitness gain

The observed changes in transcript abundance during *in vitro* fitness gain may be caused by increased gene dosage due to chromosomal amplification [8, 10]. Indeed, comparative genomic analysis of EP.1 and LP.1 parasite revealed aneuploidy for 9 chromosomes during culture adaptation, including trisomies for chromosomes (chr) 5, 23, 26, and 33, which were observed in other *in vitro* evolution experiments [8, 10] (Figure S4A and Supplementary Table 3-a and -b). We previously observed that tissue amastigotes (in infected hamster spleens) represent a mosaic karyotype, with monosomies and trisomies observed for the analyzed chromosomes, including chr 5 [10]. Based on this result, the reproducible emergence of chr 5 and chr 26 trisomies in different culture adaptation experiments represents a passive, convergent process that relies on the positive selection of pre-existing sub-populations, rather than an active, regulatory process driving karyotypic adaptation. An increased somy score was observed for these chromosomes already in EP.1, indicating a mosaic of disomic and trisomic sub-populations, the latter one showing full penetrance in LP.1. In contrast, the tetrasomy of chr 31 is stable and has been observed in all *Leishmania* species [7] and in *ex vivo L. donovani* amastigotes [8, 10]. Given the stability of this tetrasomy, regulation of expression via karyotype-dependent gene-dosage effects seems not to apply to chr 31. Thus, the expression changes between EP.1 and LP.1 observed for 144 genes are likely regulated at post-transcriptional levels (Supplementary Table 4-b). Plotting normalized genomic versus transcriptomic read depth ratios for EP.1 and LP.1 log and stat parasites correlated 75% of the up-regulated genes in LP.1 log and 42% in the LP.1 stat promastigotes with amplified chromosomes (Figure 3A, Supplementary Table 4-b and - c), affecting various biological processes associated with the LP.1 fitness tradeoff (Figure S5). Nevertheless, interrogating more specifically the read-depth ratios for trisomic chr 5 and 26 uncovered surprisingly high, gene dosage-independent fluctuations of RNA abundance in EP.1 and LP.1 promastigotes (Figure 3B and Supplementary Table 4-f and -g). While a significant fraction of transcripts on the trisomic chromosomes showed the expected 1.5-fold increase in abundance, numerous transcripts either exceeded this increase or on the contrary were expressed at lower-than-expected abundance. Such fluctuations were also observed for the LP.1/EP.1 ratios of disomic chromosomes (see chr 36, Figure 3B as an example and Supplementary Table 4-h). In contrast to the dynamic changes in karyotype, no significant fluctuations in LP.1/EP.1 read depth ratio was observed across the genome mapping the reads to 300bp genomic bins, thus ruling out episomal or intra-chromosomal amplifications as drivers of culture adaptation, at least during the first 20 passages (i.e. in LP parasites).

**Figure 3:**
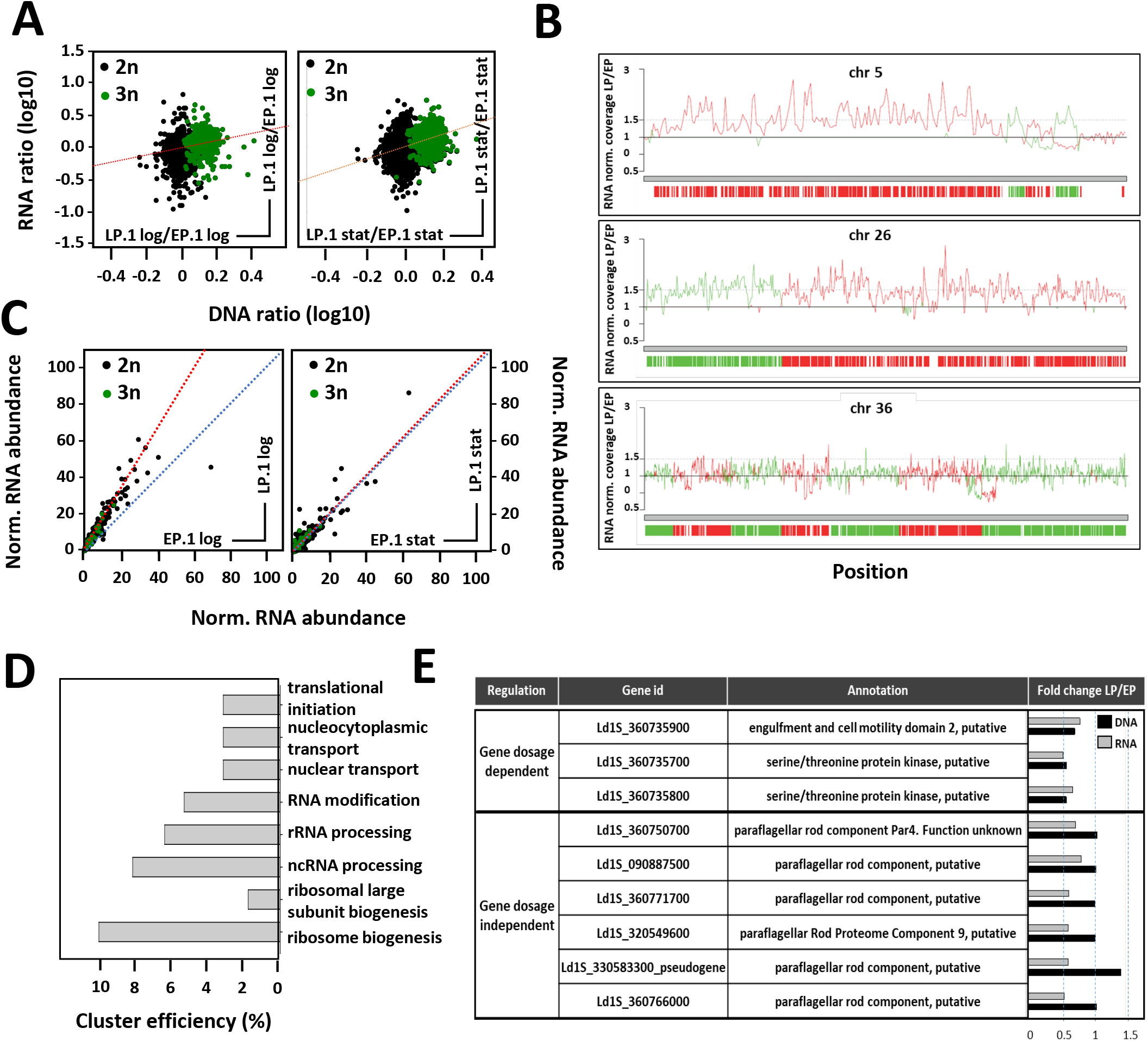
RNA abundance during fitness gain in culture is regulated by gene dosage and post-transcriptional mechanisms. (A) Ratios of DNA and RNA normalized read counts for all genes were plotted for LP.1 log compared to EP.1 log (left panel) and for LP.1 stat compared to EP.1 stat (right panel). Green dots correspond to genes encoded on trisomic chromosomes in LP.1 parasites. The regression line is represented by the dotted red line. Pearson correlation coefficients and p-values were estimated for both ratio plots using SigmaPlot software. For LP.1 log compared to EP.1 log: ρ= 0.341 and p-value < 10^-10^. For LP.1 stat compared to EP.1 stat: ρ= 0.333 and p-value < 10^-10^. (B) Normalized coverage based on the ratio of DNA read counts in LP.1 versus EP.1 for the trisomic chromosomes 5 (upper panel) and 26 (middle panel), and the disomic chromosome 36 (lower panel). The coverage ratio is indicated by the lines, while ORFs are indicated by the vertical bars. The color code reflects the DNA strand on which the ORFs are encoded (see Supplementary table 4-f to -h). (C) Post-transcriptional regulation of transcript abundance. RNA read counts were normalized by DNA read counts and plotted for all genes in LP.1 log compared to EP.1 log (left panel) and EP.1 stat compared to LP.1 stat (right panel). Green dots correspond to genes encoded on trisomic chromosomes in LP.1 (see supplementary table 4-a and -c). The calculated (red) and expected (blue) regression lines are represented. (D) Cluster efficiency computed from GO term-enrichment analysis for the ‘biological process’ category for 659 gene dosage-independent genes. Transcripts with adj. p-value < 0.01 were considered to determine the ratio of ‘normalized RNA abundance in LP.1/RNA normalized abundance in EP.1’ (see Supplementary table 4-i for details). Cluster efficiency was calculated based on 274 genes with GO IDs out of the 659 genes that showed at least a 1.2-fold increase in LP.1 normalized RNA abundance compared to EP.1. Only the functional enrichments associated with adj. p-value < 0.05 were considered. (E) Table listing selected gene dosage-dependent and -independent expression changes (from Supplementary table 4-d and -e). The fold change values computed from RNA (grey bars) and DNA (black bars) normalized read counts for LP.1 versus EP.1 log parasites are shown.

We next assessed gene-dosage independent expression changes at genome-wide level by normalizing the RNA-seq read counts to the corresponding DNA-seq reads. Direct comparison of the normalized transcript output in EP.1 versus LP.1 revealed a gene dosage-independent increase in transcript abundance for a large number of genes in LP.1 log (Figure 3C, left panel). No difference was observed for EP.1 and LP.1 stat (Figure 3C, right panel). Genome-independent, post-transcriptional increase in mRNA abundance was observed in LP.1 log parasites for genes annotated for the biological processes ‘rRNA processing’, ‘ribosome biogenesis’, ‘translational initiation’, and ‘nuclear transport’ (Figure 3D and Supplementary Table 4-i). In contrast, manual inspection revealed post-transcriptional decrease in abundance of mRNAs involved in flagellar biogenesis or EP.1-specific, ribosomal components (Figure 3E and Supplementary Table 4-j). Significantly, reduction of both DNA and RNA read depth was observed for two NIMA-related protein kinases on chr 36 that we previously associated with *in vitro* fitness gain [17] (see Figure 3E and Supplementary Table 4-d), firmly linking their depletion to accelerated growth.

In conclusion, the global analysis of the EP.1 and LP.1 transcriptomes uncovers post-transcriptional regulation as an important processes that may affect *Leishmania* fitness gain in culture, which can likely buffer against deleterious effects of genome instability and adapt mRNA abundance in a gene dosage-independent manner to a given environment.

### The fitness-adapted proteome is highly robust and enriched for GO terms associated with ribosomal biogenesis and post-transcriptional regulation

We applied a label-free, quantitative proteomics approach to assess how genomic and post-transcriptional adaptation during *in vitro* fitness gain impact protein abundance. Analyzing four independent, biological replicates of EP and LP strains (termed EP.2-5 and LP.2-5, Figure S6) identified a total of 6,050 proteins considering all samples, including 59 and 110 proteins that were exclusively detected in LP and EP parasites, respectively (Figure 4A and Supplementary Table 5-b and -f). Considering all proteins that showed a statistically different abundance (Figure 4B, Supplementary Table 5-c and -e), the majority of differentially expressed genes were shared in all four independent LP strains (566 of 788 total, 71%). These data reveal a surprising convergence of the fitness-adapted proteomes despite possible karyotypic variations between strains (Figure S4B), and inform on common pathways that are under convergent selection in LP strains during *in vitro* fitness gain. Just as observed on RNA levels, flagellar biogenesis is clearly under negative selection during culture adaptation, with reduced protein abundance observed for 46 proteins linked to flagellum and motility encoded on 24 chromosomes (Figure 4C and Supplementary Table 5-i). Another key process associated with adaptation was translation: 27 proteins encoded on 13 different chromosomes were under positive selection in LP strains (e.g. various ribosomal proteins of the 39S, 40S, 60S, L22e, and S25 families, or the ribosomal assembly protein RRB1), while only two RNA binding proteins encoded on two chromosomes were under negative selection in the same parasites (Figure 4C and Supplementary Table 5-h and -i).

**Figure 4:**
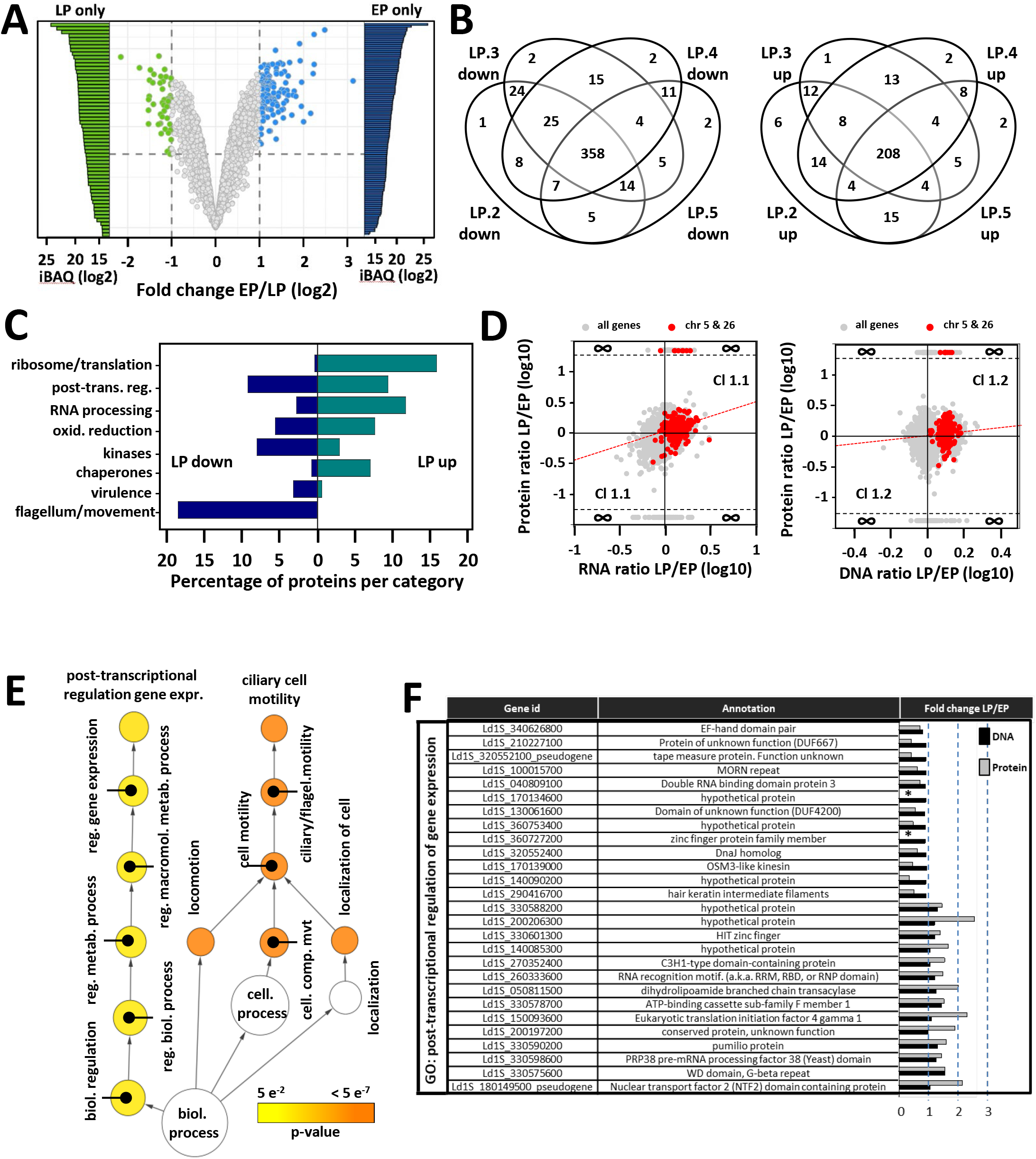
Quantitative analysis of the fitness-adapted proteome. (A) Volcano plot representing changes in protein abundance in EP log (blue dots, mean values of EP.2, EP.3, EP.4 and EP.5 are shown) compared to LP log (green dots, mean values of LP.2, LP.3, LP.4 and LP.5 are shown). Proteins identified by at least two peptides in at least three out of four biological replicates were considered. Colored dots indicate values with FDR < 0.01 and fold changes ≥ 2 (see Supplementary table 5-b and -f. The grey dots indicate non-significant expression changes. The bars indicate unique protein identifications in LP (LP only, green) and EP (EP only, blue) samples, with relative abundance indicated by the iBAQ value. (B) Venn diagram showing the number of proteins quantified and associated to a p-value < 0.01 with increased (left panel) or decreased (right panel) abundance in all four LP log biological replicates (see Supplementary table 5-c and -e. (C) Manual Gene ontology analysis of the proteins shared in all four LP log biological replicates expressed as the percentage of proteins quantified with associated p-value < 0.01 for the indicated gene categories (see Supplementary table 5-h and -i). (D) Double ratio plots comparing the fold changes computed for each gene between LP and EP log parasites for RNA (x-axis) versus protein (y-axis) (left panel) and DNA (x-axis) versus protein (y-axis) (right panel). All proteins with LFQ values were considered to determine the protein ratio LP/EP (see Methods). Grey dots represent all proteins and red dots those encoded on trisomic chromosomes 5 and 26 (see supplementary table 6-a and -h). Cluster 1.1 and 1.2 (Cl 1.1 and Cl 1.2) includes proteins whose change in abundance shows the same tendency compared to RNA abundance or gene dosage, respectively. The regression line is represented by the dotted red line. The Pearson correlation coefficients and the p-values were estimated for both ratio plots using SigmaPlot software. For protein versus RNA ratio plot: ρ= 0.349 and p-value < 10^-10^. For protein versus DNA ratio plot: ρ= 0.145 and p-value < 10^-10^. (E) Graphical representation of the GO term-enrichment analysis for the category ‘biological process’ for the 452 proteins from cluster 1 (common proteins between clusters 1.1 and 1.2), which includes 201 proteins with a GO annotation (cluster 1, see right panel D and Supplementary table 6-c). The size of the circle is indicative of the number of genes falling in each category and the color ranging from yellow to orange indicates the p-values associated as indicated in the legend. Only proteins quantified in all four biological replicates for each condition and associated with a p-value < 0.01 were considered for the GO analysis (see Supplementary table 6-b). (F) Table listing selected genes associated with the GO term ‘post-transcriptional regulation of gene expression’ from the GO enrichment analysis presented in panel E (see Supplementary table 6-c for details). Their respective fold change values computed from Protein LFQ intensities (grey bars) and DNA normalized read counts (black bars) for LP versus EP log parasites are represented. *proteins exclusive to EP log parasites.

We next assessed the level of correlation between protein abundance, gene dosage variation and transcript abundance to gain further insight into regulatory mechanisms underlying *Leishmania* fitness gain in culture. Even though the proteomics data set was obtained with four independent biological replicates (EP.2-5 and LP.2-5), the highly reproducible nature of the chr 5 and chr 26 trisomies observed in all our previous experimental evolution experiments (Figure S4B) provided a useful benchmark to assess correlations between the different data sets for at least these chromosomes. Our systems comparison suggests the presence of three different regulatory clusters for chr 5 and 26: Cluster 1 (common proteins from clusters 1.1 and 1.2) include 34 proteins whose change in abundance correlates to gene dosage and RNA abundance (Figure 4D left and right panels, upper right and lower left quadrants), including three DNA J proteins, the chaperonin 10, a HSP70 like protein and BiP, suggesting that increased stress resistance could be a potential driving force for the selection of these aneuploidies (Supplementary Table 6-j and -l). Possible post-transcriptional regulation is observed for the surface antigen-like protein (Ld1_ 050818900), whose level only correlates with mRNA abundance but not gene dosage. Finally, cluster 3 represents 5 proteins whose levels do not correlate with mRNA abundance, which are either regulated at translational levels or by protein turn-over (Supplementary Table 6-k. Thus, the increase in protein abundance is the combined result of gene dosage and mRNA abundance for the vast majority of proteins (83%) encoded on trisomic chr 5 and 26.

Gene ontology analysis of the 452 proteins that fall into regulatory cluster 1.2 (as defined by the upper right and lower left quadrants of figure 4D, right panel) revealed a strong enrichment for the term ‘post-transcriptional regulation of gene expression’ supported by 27 proteins (Figure 4E and Supplementary Table 6-b and -c), which corresponds to 15% of all proteins that show increased abundance in LP parasites (Figure S6D and Supplementary Table 6-d and -e). This enrichment is driven by the coordinated increase in expression of various proteins with known functions in RNA turnover (e.g. pumilio-domain protein encoded by Ld1S_330590200) and a series of proteins previously linked to post-transcriptional regulation in *T. brucei* such as EIF4G1 or PRP38 pre-mRNA processing factor (Supplementary Table 6-d) [56–58]. In addition, the enrichment for the GO term ‘ciliary cell motility’ is driven by the under-representation of this process in the LP proteome, supported by 20 proteins (or 18%) of all proteins showing less abundance in LP (Figure S6E and Supplementary Table 6-b, -c, -f and -g).

In conclusion, the *Leishmania* proteome undergoes reproducible, gene dosage-dependent and -independent changes during fitness gain *in vitro.* The robustness of proteomic adaptation indicates the presence of regulatory mechanisms that compensates for the genetic and transcriptomic variability between independent LP strains. At least under our experimental conditions, gene dosage-dependent changes modulate post-transcriptional regulation, which results in stabilization of various transcripts implicated in rRNA processing and ribosomal biogenesis. Thus, just as observed on transcript levels (Figures 2 and 3), the proteomic results too suggest the formation of fitness-adapted ribosomes, which in turn may control the robustness of the proteome. The role of ncRNAs in ribosomal biogenesis [59] primed us in the following to carry out a dedicated small RNome analysis in EP and LP parasites to further assess the generation of specialized ribosomes.

### Mapping the *Leishmania* non-coding transcriptome correlates snoRNA expression and rRNA modification to *Leishmania* fitness gain *in vitro*

Non-coding (nc) RNAs such as small nuclear (sn), small nucleolar (sno), ribosomal (r) or transfer (t) RNAs play essential roles in post-transcriptional regulation and translational control [24, 60]. While our data suggested an important role of these regulatory processes in genome-independent fitness gain in culture (see above), they did not inform on underlying ncRNAs as our RNAseq analyses used poly (A+)-enriched mRNA. We therefore performed a dedicated analysis of the small RNome in EP.1 and LP.1 *L. donovani* parasites. We first annotated our PacBio LD1S reference genome for ncRNAs using bioinformatics approaches (ortholog mapping, *de novo* annotation) as well as unmapped RNAseq reads of post-ribosomal supernatants (PRS) that are enriched in ncRNAs (Figure S7). These efforts established a very first repertoire of ncRNAs in any *Leishmania* species and identified 1504 genes encoding for snoRNA organized in 42 clusters on 24 chromosomes, 83 tRNA genes, 12 snRNA genes and 140 SL RNA genes (Supplementary Table 1, Figure 5A and B). Considering the trisomic chromosomes, we found 269 snoRNA genes on chr 5, 22 on chr 23, 160 on chr 26, and 193 on chr 33. We investigated more specifically the role of snoRNAs in LP.1 fitness gain in culture given the enrichment of the fitness-adapted transcriptome in the GO terms ‘ribosomal biogenesis’ and ‘rRNA processing’ (see Figure 2C). snoRNAs guide specific modifications of rRNA, such as methylation and pseudouridylation, which in turn change the specificity of the ribosome towards certain mRNAs and thus control translation [61]. We prepared whole cell lysates from both EP.1 log and LP.1 log parasites, removed the ribosomes by ultracentrifugation, and prepared libraries from the post-ribosomal supernatant (PRS). From 174 detected snoRNAs, 93 showed a more than 2-fold change in LP.1 compared to EP.1, revealing a global increase of snoRNA abundance during culture adaptation (Supplementary Table 7). Increased abundance was confirmed for 7 out of the 8 snoRNAs probed by Northern blot analysis of the PRS (Figure 5C and D).

**Figure 5:**
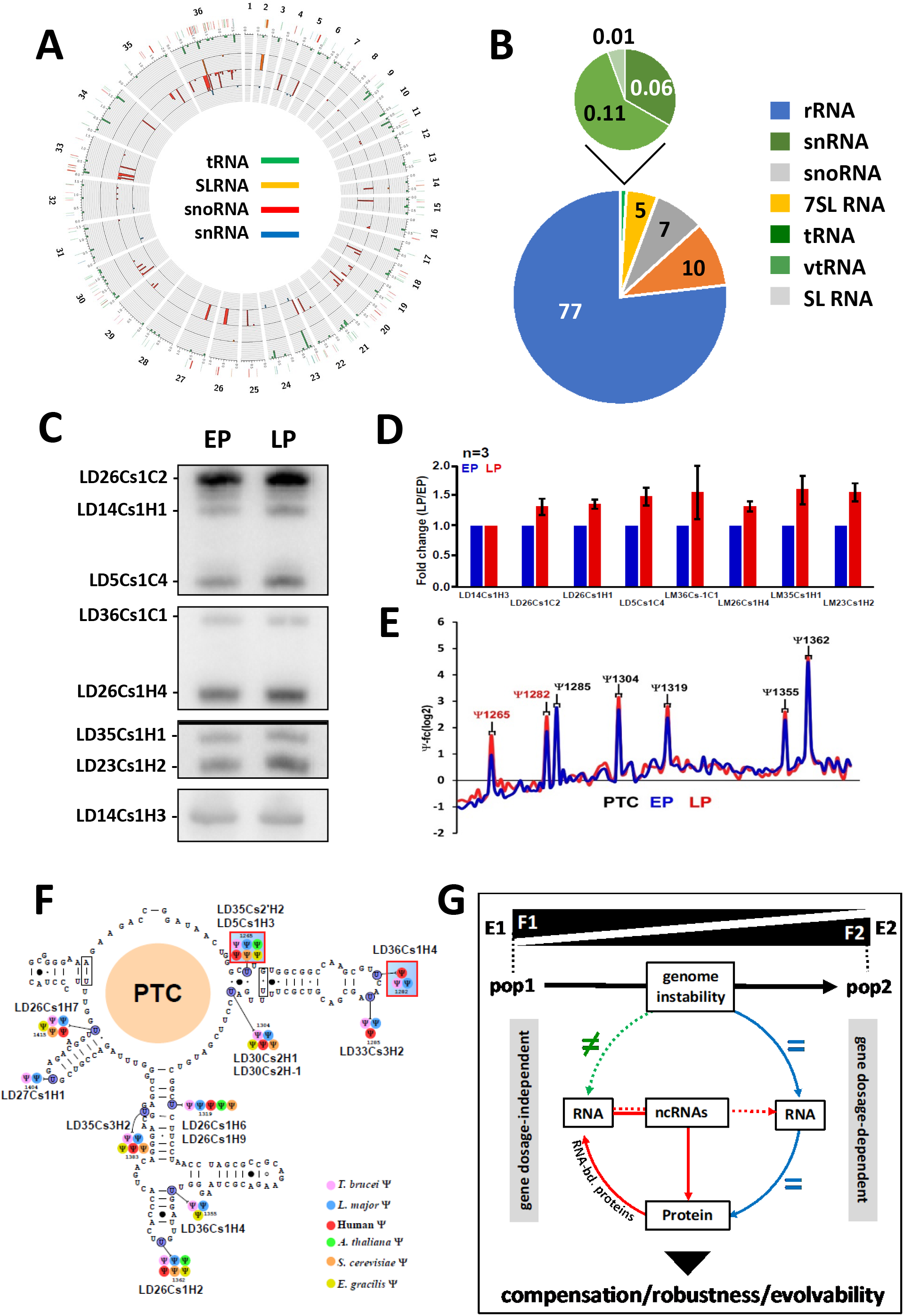
The fitness tradeoff in LP promastigotes correlates with snoRNA expression changes and increased rRNA pseudouridinylation levels. (A) Genomic map of *L. donovani* Ld1S ncRNA genes. (B) Composition of the small RNome identified in EP parasites. (C) Northern blot analysis of selected snoRNAs, Ld14Cs1H3 was used as loading control. One representative northern blot out of three is presented. (D) Histogram plot representing the fold changes between LP (red bars) and EP (blue bars) log parasites corresponding to densitometric analysis of the signals shown in (C). (E) Representative line graph of the fold change in rRNA pseudouridinylation level (Ψ-fc, log2) is presented for EP (blue line) and LP (red line). The positions where the Ψ level is increased in three replicates are indicated in red. (F) The location of Ψ sites in the rRNA is depicted on the secondary structure. Hypermodified sites are highlighted in red squares. The snoRNAs guiding each Ψ are indicated. The color code for each Ψ site is indicative of the organism where it was already reported. (G) Model of *Leishmania* evolutionary adaptation. Different environments (E1, E2) select for different fitness traits (F1, F2), which modify the parasite population structure (pop 1, pop 2). In the absence of transcriptional regulation, *Leishmania* exploits genome instability to generate changes in gene dosage via chromosome and gene copy number variations. These changes are either correlated (blue arrows) or not (green arrow) to changes in transcript and protein abundance. The gene dosage-regulated transcriptome and proteome (right panel) is highly enriched for the GO term ‘post-transcriptional regulation of gene expression’ and thus likely regulates gene dosage-independent changes in RNA abundance (red arrow, left panel). The enrichment of these transcripts in ncRNAs in turn can control RNA stability and translatability by guiding modifications of mRNA or rRNAs. This allows for (i) compensation of deleterious gene dosage effects, (ii) phenotypic robustness despite genetic heterogeneity, and (iii) maintenance of evolvability despite selection pressure.

Next, we examined if the increase in snoRNA abundance affected the level of rRNA pseudouridylation (Ψ) by applying a modified RNAseq protocol (termed Ψ-seq) using total RNA from EP.1 and LP.1. We detected two hyper-modified rRNA sites in all three biological replicates at positions Ψ1265 and Ψ1282 inside the peptidyl transferase center (PTC) (Figure 5E), which correlated with the increased abundance of the corresponding snoRNAs that guide these modifications (Figure 5F, Supplementary Table 7) [62]. Our data thus provide a first link of snoRNA expression and rRNA modification to *Leishmania* fitness gain in *in vitro* culture, which further supports the possibility of fitness-adapted ribosomes and suggests translational control – in addition to genomic and post-transcriptional adaptation – as yet another, gene dosage-independent mechanism likely linked to *Leishmania* evolutionary adaptation.

## Discussion

A common hallmark of all microbial pathogens is their capacity to adapt to unpredictable fluctuations in their host environments through an evolutionary process, where genetically heterogenous individuals constantly compete for survival [63]. Here we combined experimental evolution and integrative systems approaches to uncover mechanisms of fitness gain in the human pathogen *Leishmania donovani*. Our study provides new evidence that these parasites combine regulatory processes at genomic, post-transcriptomic and translational levels to establish highly robust fitness phenotypes while maintaining genetic heterogeneity thereby avoiding genetic death.

In the absence of classical, promoter-driven control of gene expression, *Leishmania* relies on alternative mechanisms to regulate transcript and protein abundance, including regulated mRNA turn over and translational control [5, 64]. These parasites further use a highly unusual, genomic form of gene expression regulation, where changes in chromosome and gene copy number control transcript abundance via gene dosage [7–10]. Previous studies allowed us to link these forms of genome instability to fitness gain *in vitro* as judged by the highly reproducible karyotypic changes observed during culture adaptation in independent clinical and animal-derived *L. donovani* isolates [8, 10, 20]. Positive selection of chromosome amplification is further sustained by the independent evolutionary experiments conducted in this study, which once more revealed amplification of chromosomes 5 and 26 as key drivers of *in vitro* fitness gain. Such karyotypic changes are not exclusive to culture adaptation but have been documented in *L. donovani* tissue amastigotes [10], and in drug resistant clinical *Leishmania* isolates [13]. Similar to stress-adaptation in fungi [65], karyotypic changes thus may provide the genetic diversity required for *Leishmania* to evolve beneficial phenotypes in response to environmental change. However, such structural mutations simultaneously affect the expression level of hundreds of genes, raising questions on the nature of the coding sequences that drive karyotypic selection during parasite adaptation, and on the mechanisms that suppress deleterious gene dosage effects while preserving beneficial ones. Applying an integrative systems approach on promastigote parasites during culture adaptation (early passage, EP and late passage, LP) allowed us to address these important open questions.

Comparative genomic, transcriptomic and proteomic analyses of EP and LP promastigote parasites revealed a gene dosage-dependent increase in mRNA and protein abundance for genes implicated in RNA turnover, including RNA-binding proteins known to regulate mRNA stability [66], pumillo domain proteins known to regulate ncRNA abundance [67], and a series of proteins that were associated with trypanosomatid mRNA-binding and post-translational regulation in recent, genome-wide functional screens [56–58]. Positive selection of chromosome amplifications during *L. donovani* culture adaptation thus is likely driven by genes that establish an adaptive, post-transcriptional interface. This interface may regulate differential mRNA stability during fitness gain in culture, which can compensate for deleterious gene dosage effects by selective mRNA degradation, while at the same time boosting the expression of beneficial genes conferring stability to selected mRNAs. Assessment of gene-dosage-independent expression changes indeed correlated both increased and decreased mRNA abundance to the observed fitness phenotype. In the absence of transcriptional control in *Leishmania*, these gene dosage-independent changes in mRNA abundance must be regulated by differential RNA stability. A number of transcripts implicated in flagellar biogenesis showed reduced stability during culture adaptation, which correlated with reduced mobility (data not shown). This coordinated process likely involves shared cis-regulatory sequence elements in the transcripts’ 3’UTR that are recognized by the RNA-binding proteins [5]. Loss of flagellar function associated to *L. donovani in vitro* fitness gain has been observed in other independent evolutionary experiments [17] and thus represents a reproducible, convergent phenomenon that may liberate ATP for energetically demanding processes that are under positive selection during culture adaptation. Indeed, a large number of transcripts implicated in highly energy-demanding ribosomal biogenesis and translation were stabilized in LP parasites. The differential regulation of mRNA abundance observed in our experimental evolution system thus establishes a first link of post-transcriptional regulation to *Leishmania* fitness gain in our *in vitro* setting.

In culture, fitness (defined as reproductive success of a given population) is largely synonymous to proliferation, which depends on the number of ribosomes and the cell’s translational potential [68]. While the fitness-adapted transcriptome is indeed characterized by increased expression in ribosomal components thus fueling the need for more ribosomes, the differential expression of various 40S and 60S ribosomal protein isoforms in LP compared to EP parasites further suggests that adaptation is linked to a qualitative, ribosomal changes (see Supplementary Tables 2 and 5). Such dynamic regulation of ribosomal biogenesis may give rise to specialized ribosomes, which not only may increase translation efficiency in these fast-growing LP parasites, but could also control translation of unwanted mRNAs, thus providing an additional filter (next to differential RNA stability) to eliminate toxic gene dosage effects. The existence of such structurally distinct, specialized ribosomes has been observed in *Plasmodium* spp., where stage-specific expression of certain rRNA isoforms allows for the establishment of A-type (asexual stage specific) and S-type (sporozoite specific) ribosomes [69, 70]. Likewise, stage-specific modification of rRNA has been linked to the transition of *T. brucei* from the procyclic insect to the mammalian bloodstream forms [23]. Finally, changes in expression and modification of different rRNA genes, ribosomal proteins, and translation factors indeed can control preferential translation of different subsets of mRNAs in other organisms [71], including *Dictyostelium discoideum* [72, 73], zebrafish development [74], or cancer [75].

Conducting a dedicated analysis of the *L. donovani* non-coding (nc) RNome, we have provided further support to the existence of such fitness-adapted ribosomes in *Leishmania*. First, we observed post-transcriptional upregulation of a large number of snoRNAs and five (out of a total of 9) pseudouridylate synthases in LP compared to EP promastigotes (see Supplementary Table 4). Second, these snoRNA expression changes correlated to changes in the pseudouridinylation (Ψ) profile of the rRNA peptidyl transferase center (PTC) that catalyzes peptide bond formation and peptide release [76]. Similar Ψ hyper-modification of rRNA was previously observed in bloodstream form trypanosomes and likely contributes to stage-specific adaptation [23]. Given the high coding density of chr 5 and chr 26 for snoRNAs and the functional enrichment of these chromosomes for GO term ‘rRNA processing’ (see Figure S4C), it is interesting to speculate that their convergent amplification in all our evolution experiments may be driven by their ncRNA content and their requirement for fitness-adapted translation. Indeed, defects in rRNA pseudouridylation affect ribosomal ligand binding and translational fidelity in eukaryotic cells [77], and changes in PTC modification were shown to affect both the ribosome structure and activity in yeast [78]. The combination of (i) different rRNA isoforms, (ii) hundreds of snoRNAs and differentially modified rRNA sites, (iii) diverse 40S and 60S ribosomal proteins, and (iv) the formation of different translation complexes [79–81] defines a vast ribosomal landscape in *Leishmania.* Translational control via fitness-adapted ribosomes likely fine-tunes expression and provides proteomic and phenotypic robustness to adapting parasite populations, which thus can maintain genetic diversity and evolvability despite constant natural selection [10].

In conclusion, our data uncover *Leishmania* evolutionary adaptation as an emergent property of a highly complex process that integrates variations in gene dosage with correlating changes in transcript abundance for genes implicated in post-transcriptional regulation and ribosomal biogenesis, which may compensate for toxic gene dosage effects via differential RNA turn-over and translational regulation, respectively (see Figure 5G). Even though our results are largely correlative in nature, our model is supported by the convergence of the genomic, transcriptomic and proteomic signals we observed between independent populations, which thus are the result of natural selection rather than random genetic drift. Our findings challenge the current genome-centric approach to *Leishmania* epidemiology and suggest the *Leishmania* non-coding RNome as well as regulatory circuits at transcriptional and translational levels as potential novel sources for biomarker discovery in clinical settings. Finally, our model may be of relevance to other pathogenic systems that gain fitness through genome instability, including fungal infection and cancer.

## Supporting information

Supplementary Table 2bis

## Acknowledgements

This work was supported by the Fondation de la Recherche Médicale contract FDT201805005619 (LP), the Agence Nationale pour la Recherche Labex ‘Integrative Biology of Emerging Infectious Diseases’ contract ANR-10-LABX-62-IBEID and Labex ‘French Alliance for Parasitology and Health Care’ contract ANR-11-LABX-0024 (GFS, LP, PP, GB), the Campus France Franco-Israeli Programme Hubert Curien Maimonide 2018 (GFS and SM), the seeding grant from the Institut Pasteur International Department to the LeiSHield Consortium and the EU H2020 project LeiSHield-MATI - REP-778298-1 (GB), the France Génomique National infrastructure, funded as part of the « Investissements d’Avenir » program managed by the Agence Nationale pour la Recherche contract ANR-10-INBS-09 (HV, RL, CP), and the ERD Funds, project CePaViP (CZ.02.1.01/0.0/0.0/ 16_019/0000759) (BV, JS, PV). We thank the CEA-CNRGH for its contribution to the sequencing costs and all the CEA-CNRGH staff who contributed to sample preparation and sequencing for their excellent technical assistance.

The authors declare that they have no conflict of interest.

## Supplementary Figures

**Figure S1:**
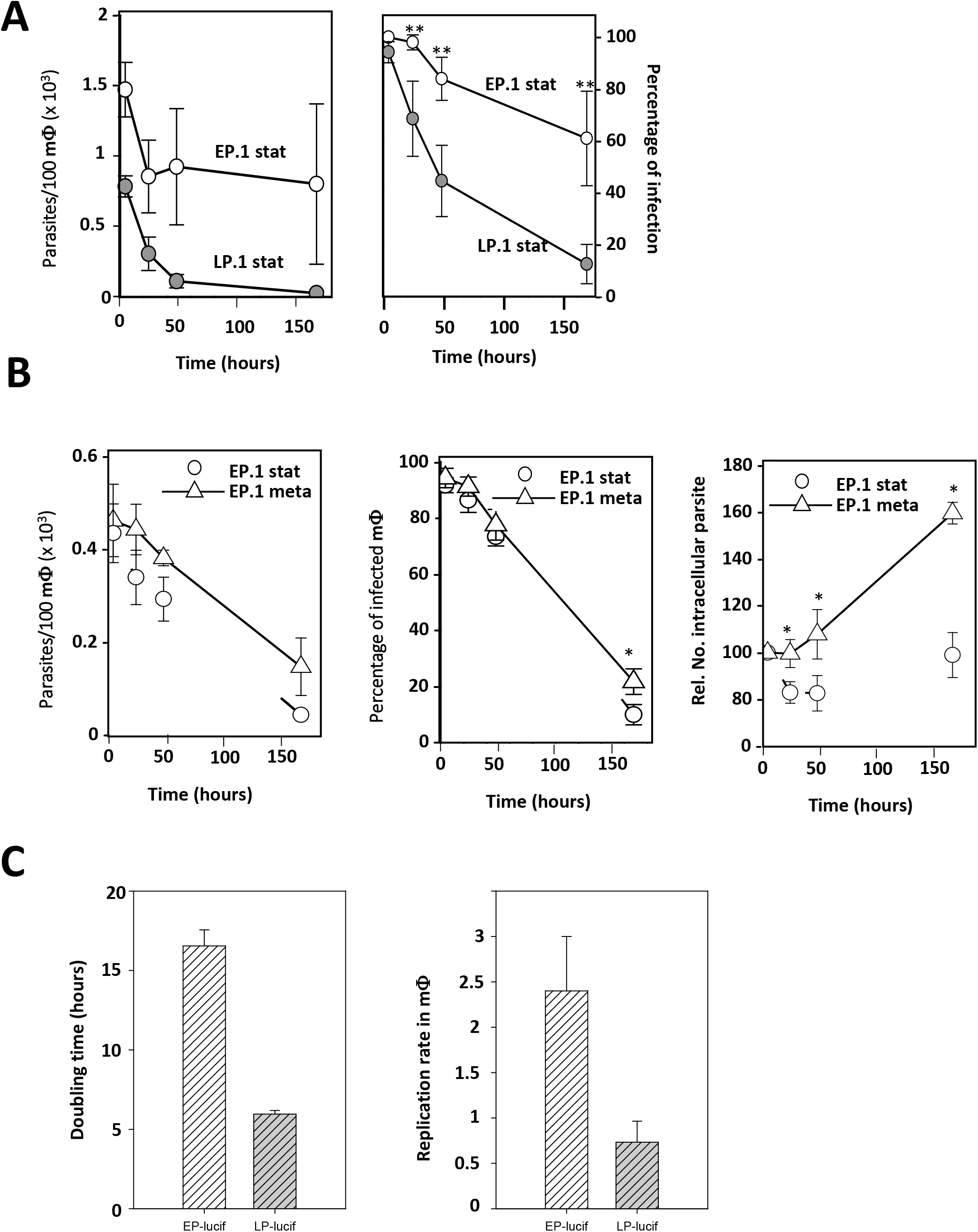
Parasite growth and macrophage infection studies. (A) Comparison of EP.1 stat (open circle) and LP.1 stat (grey circles) infectivity. Mean number of parasites per 100 macrophages +/-SD (left panel) and the percentage of infection (right panel) from three biological replicates are shown. (B) Comparison of EP.1 stat (open circle) and EP.1 meta (open triangle) infectivity. Mean number +/-SD of parasites per 100 macrophages (left panel), the percentage of infection (middle panel) and the relative number of intracellular parasites (right panel) from a representative experiment out of three replicates are shown. * indicates p-value ≤ 0.05. (C) Histogram plots representing the generation time of EP and LP parasites originated from an independent evolutionary experiment with parasites expressing luciferase (EP.luc and LP.luc) (left panel). Replication rate in infected macrophages for the EP.luc and LP.luc parasites calculated between day 1 and day 6 after infection (right panel).

**Figure S2:**
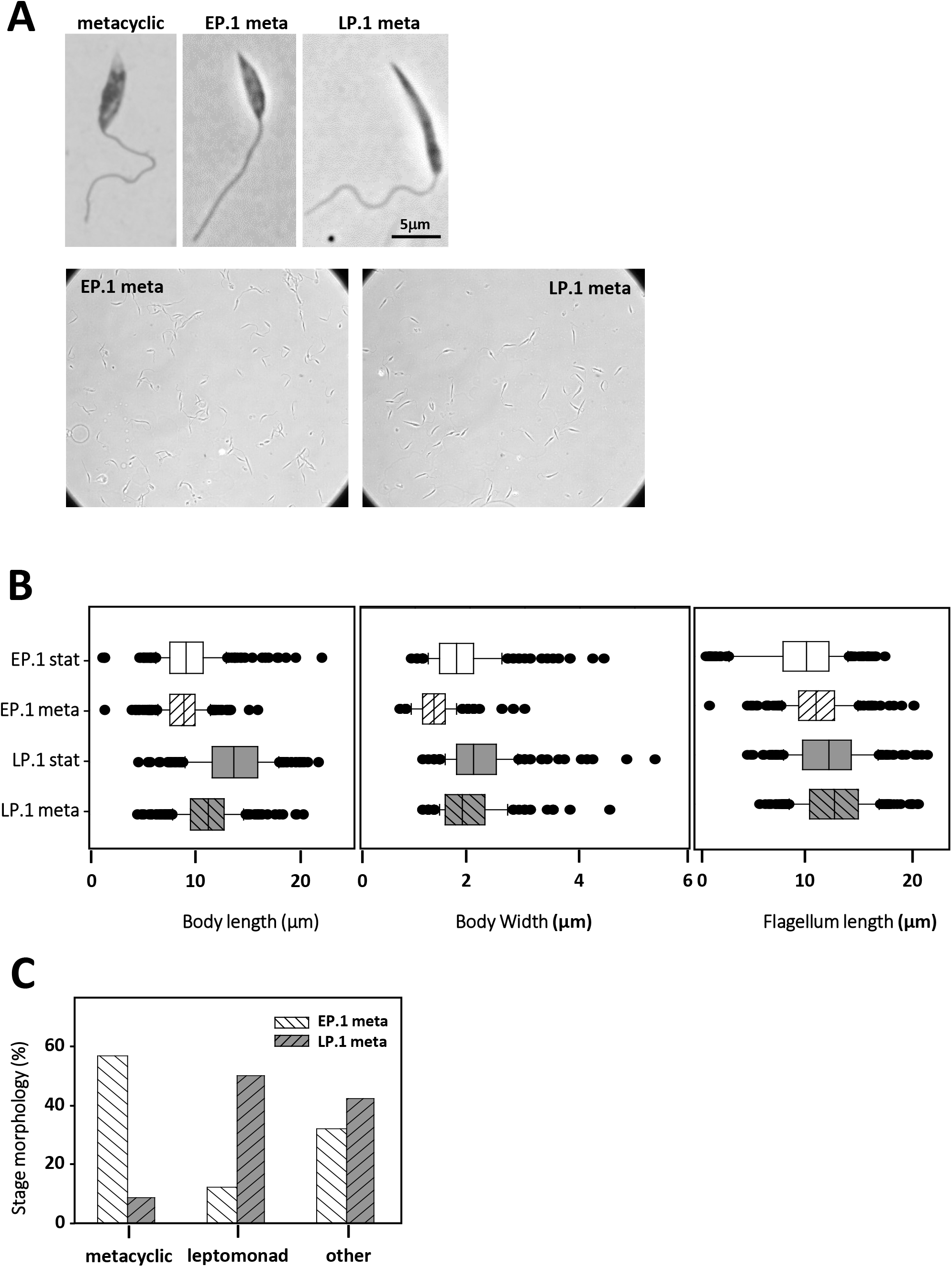
Morphological analysis of EP and LP metacyclic-enriched parasite fractions. (A) Micrographs of representative EP.1 metacyclic isolated from the sand fly thoracic part (upper left image), Ficoll-enriched EP.1 (upper middle image) and LP.1 metacyclic-like parasites (upper right image) from stationary culture. Broad field images of EP.1 and LP.1 metacyclic enriched parasites are presented in the lower right and left images. (B) Quantitative morphological analysis of stationary-phase and metacyclic-enriched parasite populations. The box plots show the median values and the upper and lower quartiles for body length (left panel), body width (middle panel) and flagellum length (right panel). (C) Distribution of the indicated promastigote forms in EP.1 and LP.1 Ficoll-enriched metacyclic fractions.

**Figure S3:**
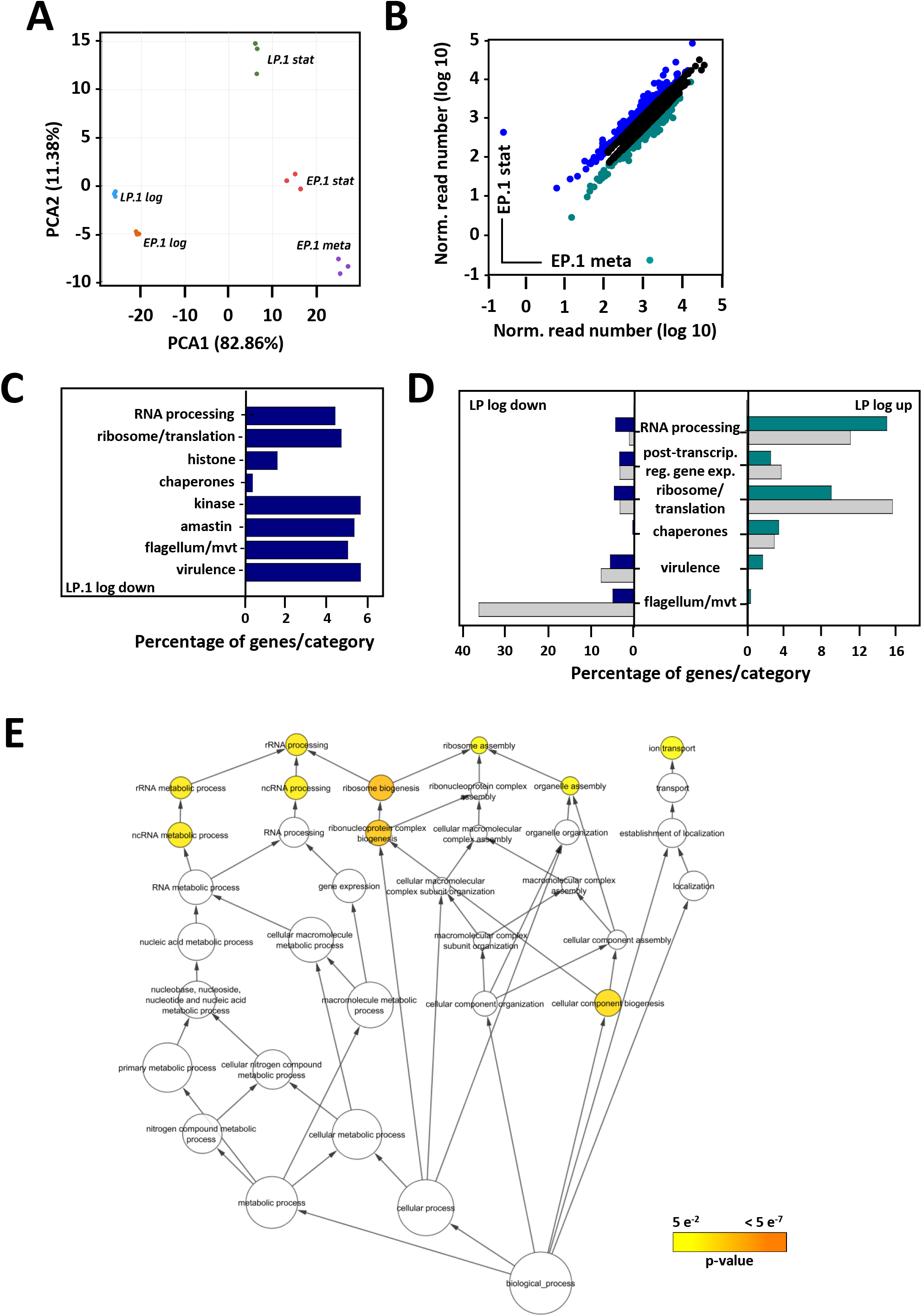
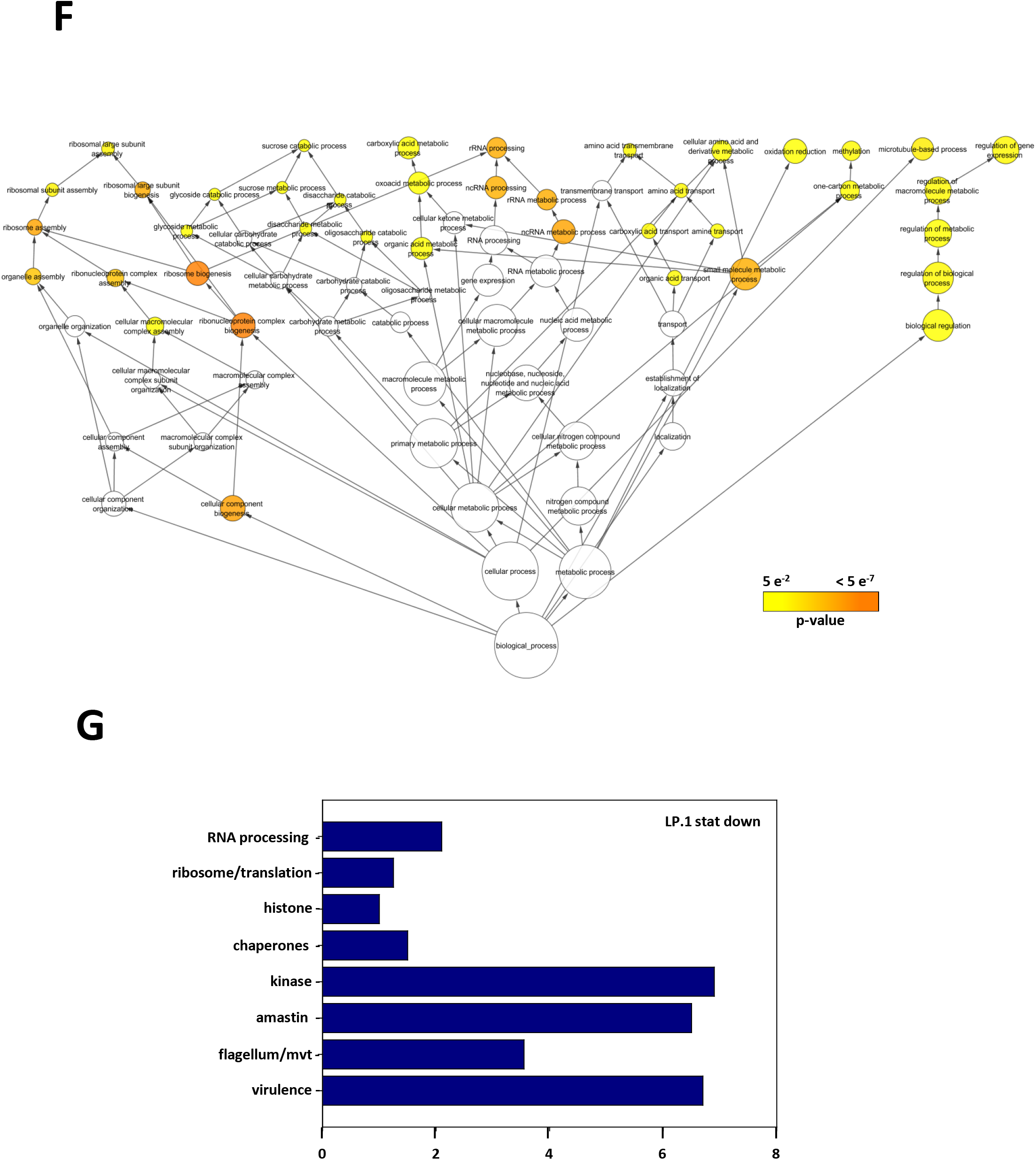
Transcriptomic analysis of EP.1 and LP.1 parasites. (A) Principal component analysis of EP.1 and LP.1 parasites from logarithmic (log) and stationary (stat) phase cultures, and after metacyclic enrichment (meta). (B) Ratio plots of normalized RNA abundance for EP.1 stat compared to EP.1 meta. Dark blue and dark cyan dots represent respectively gene expression changes with FC > 1.5 and adjusted p-value ≤ 0.01; black dots correspond to gene expression changes with adjusted p-value > 0.01. (C) Histogram plot representing the number of genes showing decreased transcript abundance in LP.1 log for the indicated gene categories (see supplementary table 2-j). (D) Histogram plot showing the number of genes with decreased and increased transcript abundance in LP log parasites from two independent transcriptomic analyses. Cyan and blue histogram bars represent the evolutionary experiment presented in Figure 2 (EP.1 and LP.1), grey bars correspond to the second RNAseq data set corresponding to EP.8, EP.9, LP.9 and LP.10 samples (see Figure S8 and Supplementary table 2bis-c and -e for detail). (E, F) Graphical representations generated with the BiNGO plugin of the Cytoscape software package for the GO term-enrichment analysis performed with the transcripts showing statistically significant increased abundance in LP.1 log (E) and LP.1 stat (F) (see Supplementary table 2-o). The size of the circle is indicative of the number of genes falling in each category and the color ranging from yellow to orange indicates the p-values associated as indicated in the legend. (G) Histogram plot representing the number of genes showing decreased transcript abundance in LP.1 stat for the indicated gene categories (see supplementary table 2-n).

**Figure S4:**
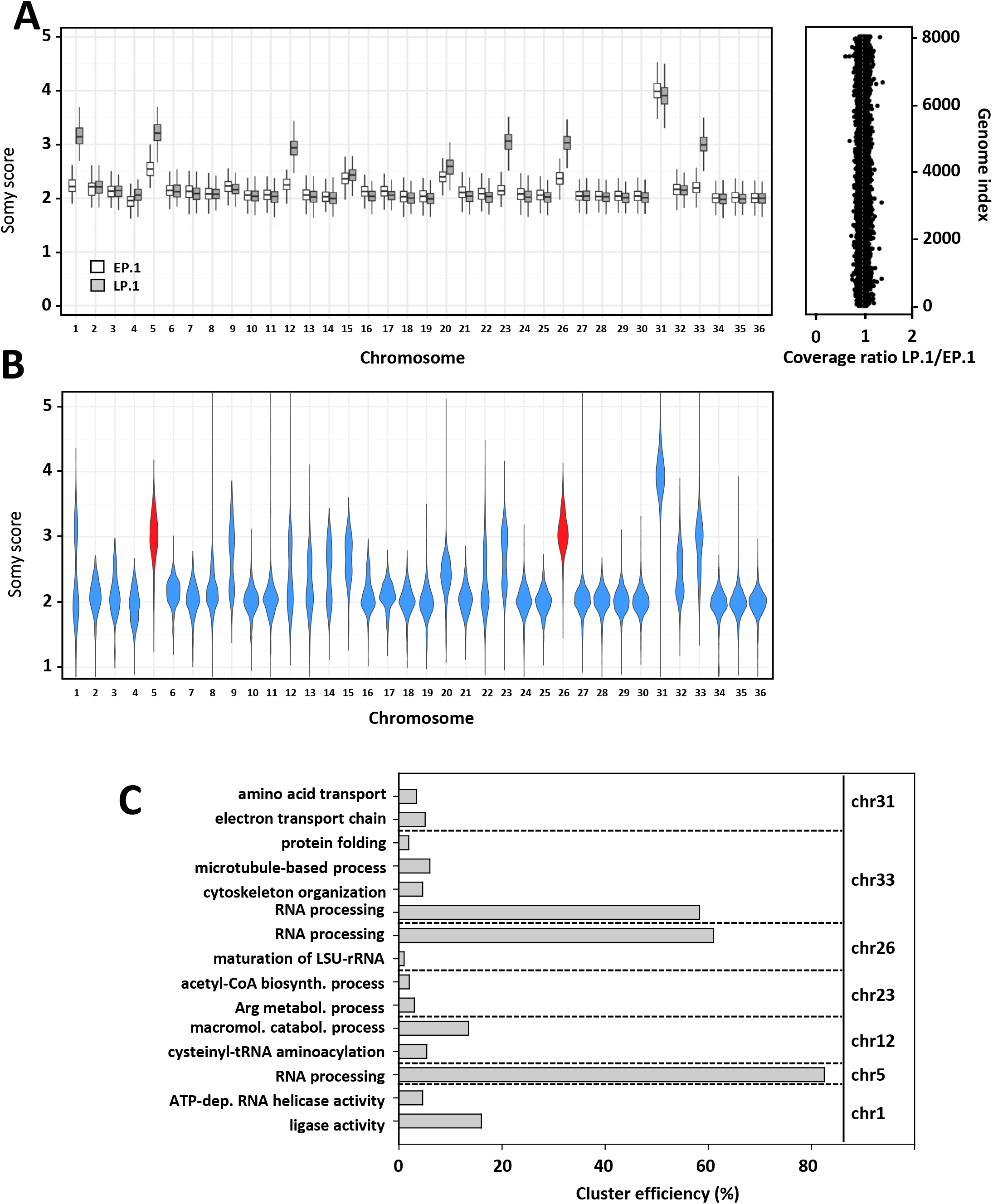
Comparative genome analysis of EP and LP parasites from independent evolutionary experiments. (A) Chromosome somy levels of EP.1 and LP.1 promastigotes. Chromosome read-depth distributions are shown in boxplots depicting the median and the upper and lower quartiles (left panel). Genome-wide coverage ratios (y axes) between LP.1 and EP.1 (right panel). Genome-wide coverage ratio (x-axis) between EP.1 and LP.1. The y axis reports the position of the genomic windows along the chromosomes. Dots represent genomic windows of 300 bases. (B) Violin plot computed from three independent evolutionary experiments representing the somy score distribution for each chromosome. In red are highlighted chr 5 and 26 that are trisomic in all three experiments (LP.1, LP.6 and LP.7). (C) Enrichment analysis of the aneuploid chromosomes for the GO categories ‘molecular function’ (chr 1) and ’biological process’ (5, 12, 23, 26, 31 and 33). The bars correspond to the cluster efficiency computed from GO term-enrichment analyses (see Supplementary table 3-d).

**Figure S5:**
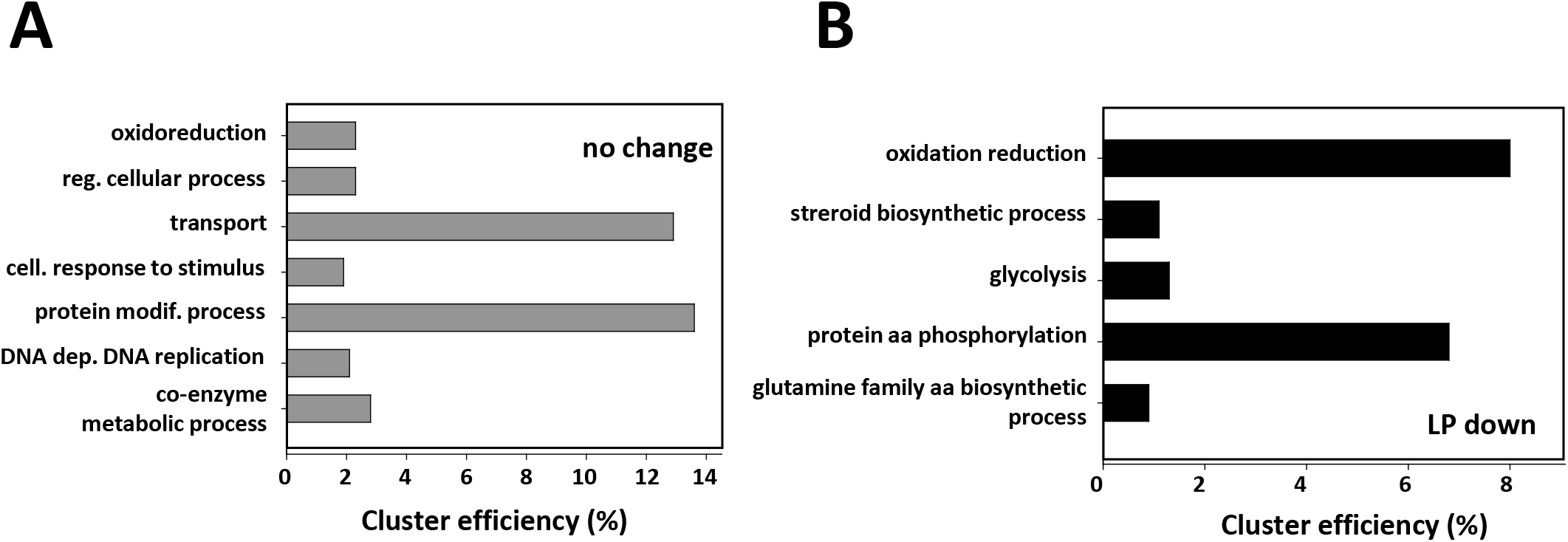
GO analysis of gene dosage-dependent and -independent changes in RNA abundance. (A, B) Enrichment analysis for the GO category ’biological process’. RNA read counts were first normalized by DNA read counts to estimate the ratio of normalized RNA abundance between LP.1 and EP.1 (see Supplementary table 4-i). (A) Histogram showing the cluster efficiency for 1104 genes that show dosage dependent changes in mRNA abundance (ratio from 0.8 to 1.2), including 463 genes that are annotated with a GO term. (B) Histogram showing the cluster efficiency for 1192 genes that show dosage in-dependent changes in mRNA abundance (ratio < 0.8), including 510 genes that are annotated with a GO term and show a decrease in RNA read counts after normalization to DNA read counts in LP.1 log parasites.

**Figure S6:**
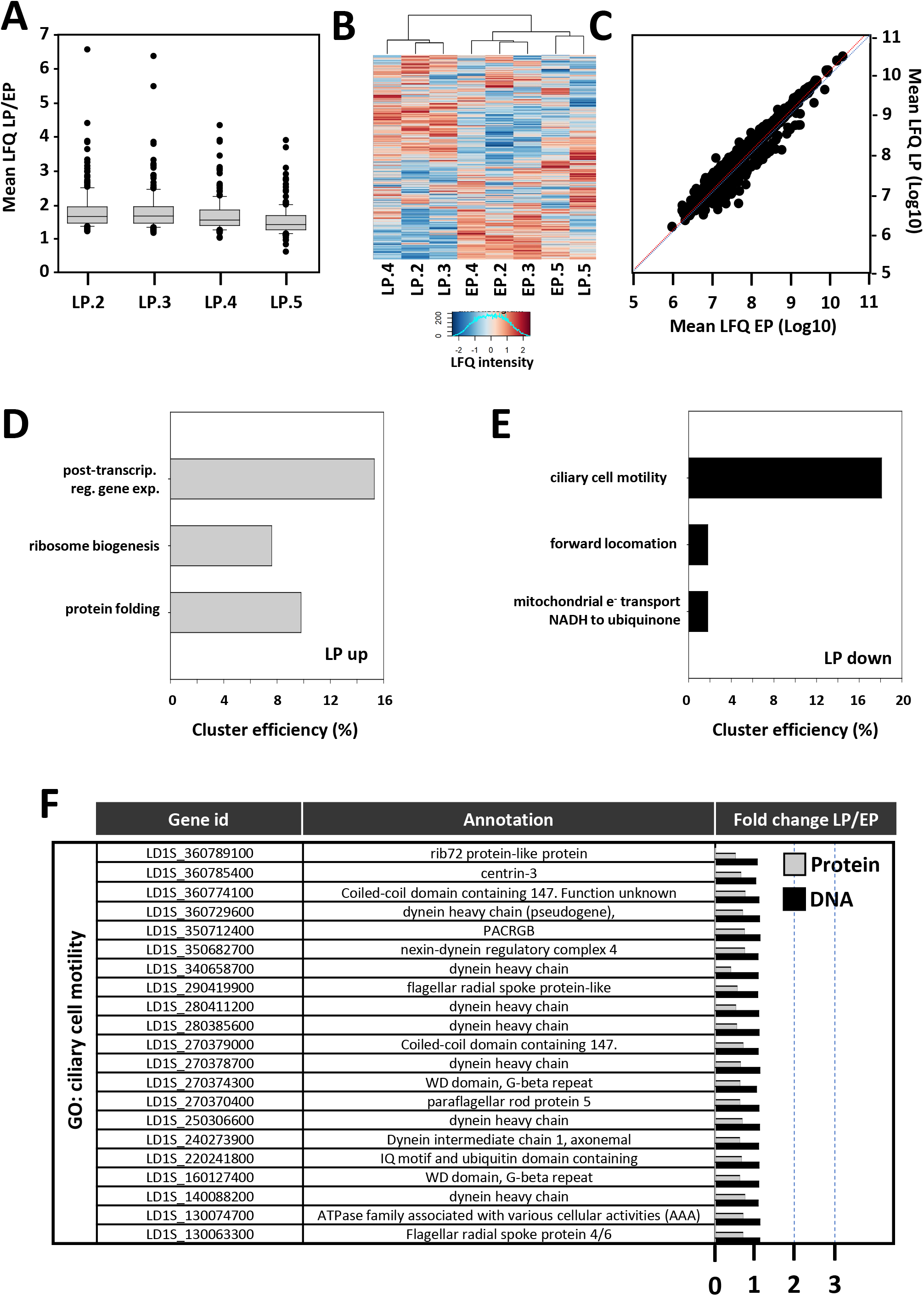
Quantitative proteomics analysis. (A) Box plots representing the median ratio and the upper and lower quartiles of the LFQ intensity values for all LP biological replicates (LP.2, LP.3, LP.4 and LP.5) compared to the median of all EP replicates (EP.2, EP.3, EP.4 and EP.5). (B) Cluster analysis of all EP and LP samples (Ward method). (C) Ratio plot representing the mean LFQ intensity value between EP and LP for each individual, quantified protein. The experimental and the expected regression lines are shown in red and blue respectively. (D, E) Cluster efficiency for the GO category ‘biological process’ for proteins from cluster 1 whose abundance correlates with increased (D) or decreased normalized DNA read counts (E) in LP log parasites. Only proteins quantified with a p-value < 0.01 were considered for the GO term enrichment analysis (see Supplementary table 6-e and -g). (F) Table listing selected genes associated with the GO term ‘ciliary cell motility’ from the GO enrichment analysis presented in Figure 4E. Their respective fold change values computed from Protein LFQ intensities (grey bars) and DNA normalized read counts (black bars) for LP versus EP log parasites are represented.

**Figure S7:**
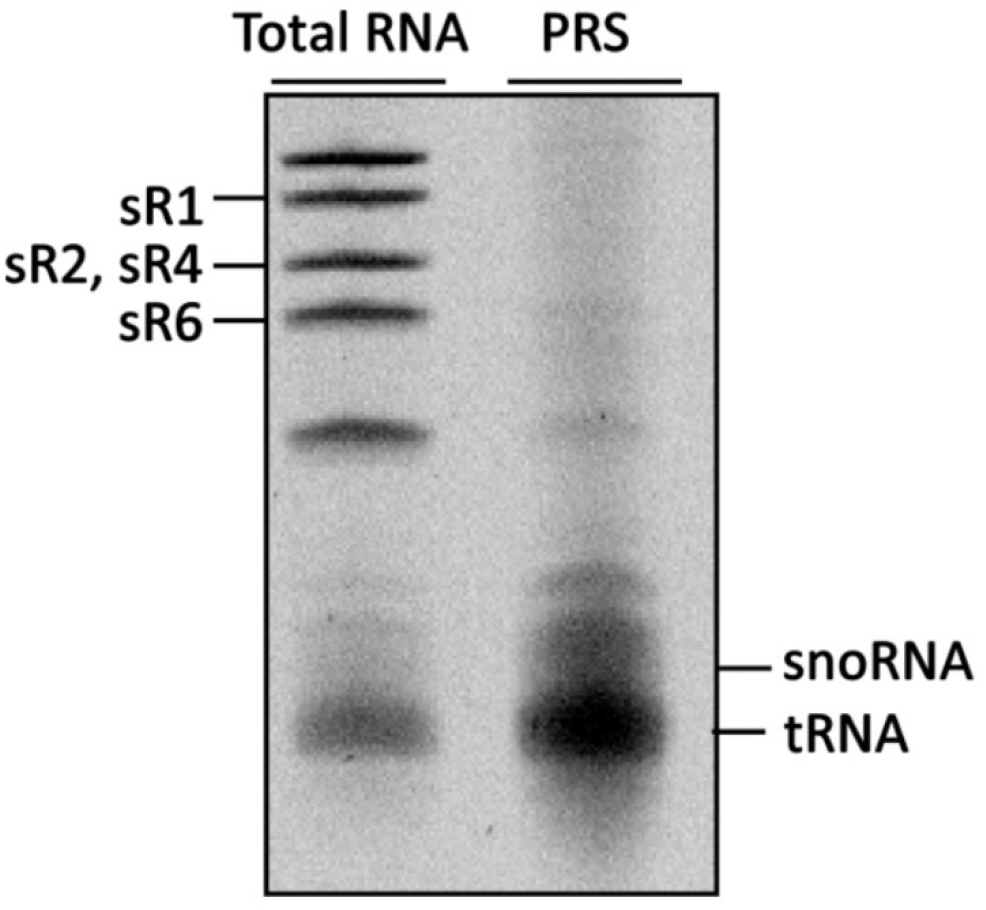
Enrichment of small RNAs obtained from post-ribosomal supernatants (PRS) of EP promastigotes. 2×10^9^ cells were disrupted by nitrogen cavitation under low salt concentration (150 mM KCL) in the presence of high MgCl_2_ (10 mM) followed by ribosome extraction using high KCL (300 mM). The ribosomes were removed by centrifugation at 35,000 rpm for 2h. 2µg of RNA from total lysate (Total RNA) and PRS sample were separated on a 10% polyacrylamide gel and stained with ethidium bromide.

**Figure S8:**
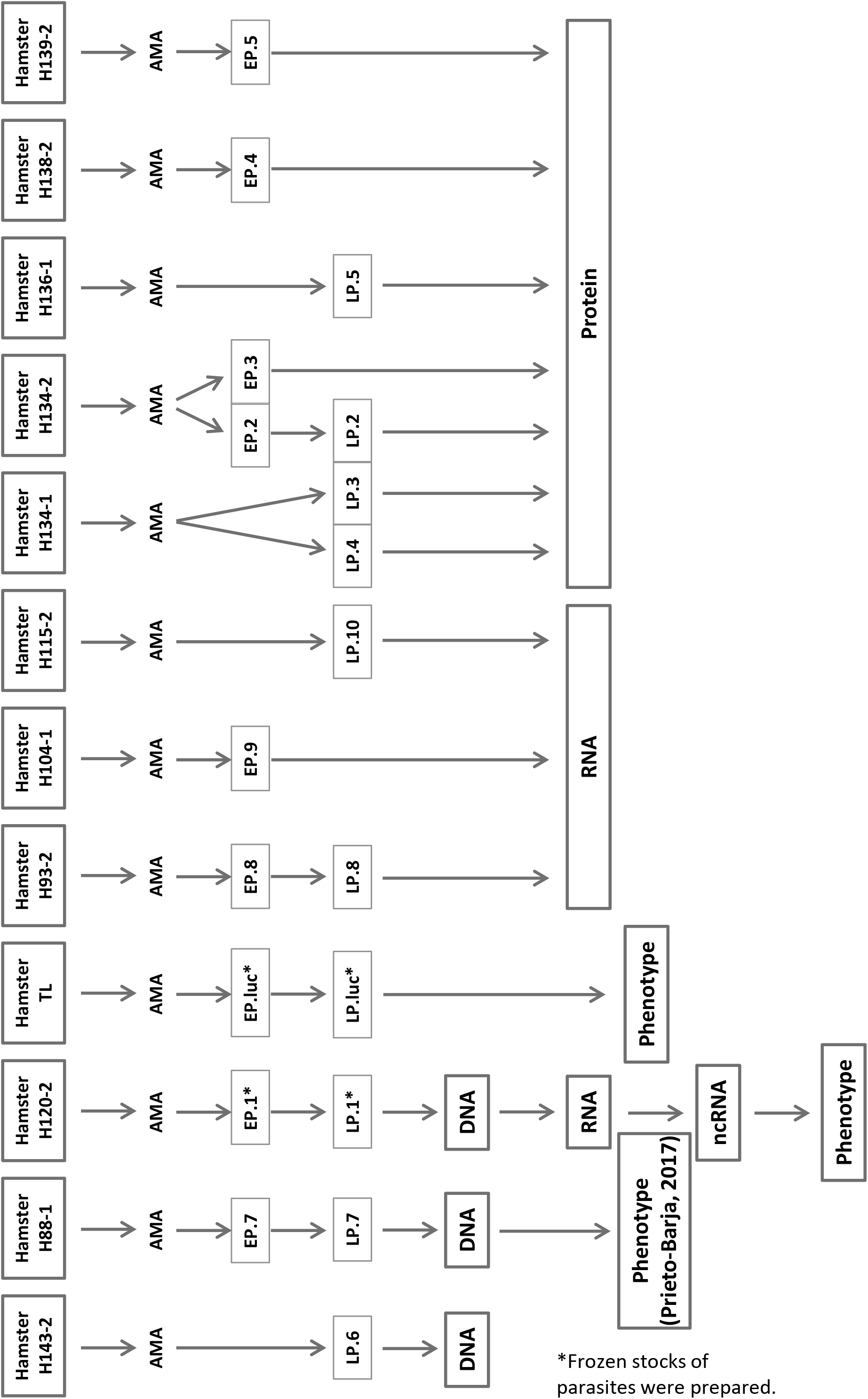
Overview chart of strains used in this study. Each hamster infected with *L. donovani* parasites was identified by the cage number and is the source of amastigotes (AMA) for conversion to promastigotes. Early passage promastigotes (EP) and late passage promastigotes (LP) used for the genomic (DNA), transcriptomic (RNA), small RNome and transcriptome-wide mapping of pseudouridine sites (RNome & ᴪ-seq), proteomic (Protein) and phenotypic analyses are identified. The parasites marked by an asterisk (*) were frozen at passage 2 and passage 20.

**Figure S9:**
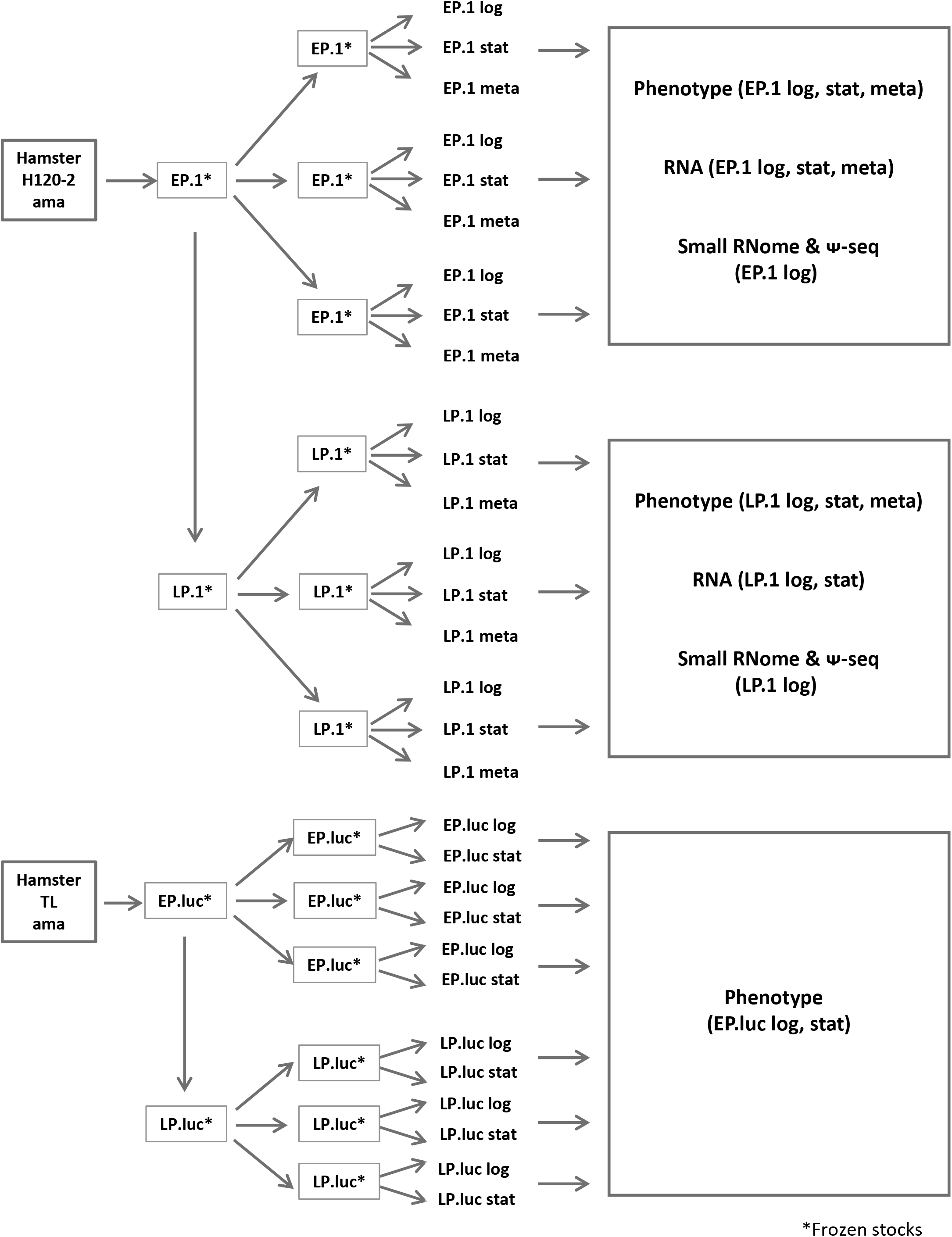
Experimental flow chart. Strains issued from independent experimental evolution assays are identified by number (i.e. EP.1 and LP.1 are the strains resulting from experiment 1) (see Figure S8 for details). Frozen stocks of EP.1, LP.1, EP.luc and LP.luc were prepared. The stage-specific expression analysis was therefore performed starting from three frozen aliquots prepared at passage 2 (EP.1) and passage 20 (LP.1). Each of the frozen parasites was used to prepare RNA extracts from log and stationary growth culture and from enriched metacyclic forms (see Figure S9). Likewise, phenotypic analyses performed with EP.1, LP.1, EP.luc and LP.luc started from frozen aliquots for each replicate (see Figure S9).

## Supplementary Tables

Supplementary Table 1. Transcriptomic read counts of EP and LP RNAseq analyses.

Supplementary Table 2. EP.1 versus LP.1 comparative transcript profiling.

Supplementary Table 2bis. EP versus LP comparative transcript profiling (EP.8, EP.9, LP.8 and LP.10.

Supplementary Table 3. Genomic analyses of EP and LP parasites.

Supplementary Table 4. Analysis of gene copy number-independent changes in RNA abundance.

Supplementary Table 5. Proteomic analysis of EP and LP log parasites.

Supplementary Table 6. Correlation between protein abundance, gene dosage variation and transcript abundance.

Supplementary Table 7. snoRNA expression levels in EP.1 and LP.1 parasites.

