## Supplementary Table 7 for "Experimental evolution reveals post-transcriptional regulation as a novel driver of *Leishmania* fitness gain"

**Supplementary Table 7**. **snoRNA expression level in EP and LP parasites.** RNA from both type pf parasites was used to prepared PRS RNA. The number of the read are given in Reads per kilo million (RPKM). Fold change (FC) is given for RPKM of (LP/EP) for both replicates. The identity, length in nt, chromosomal location, and the type of snoRNAs (H/ACA or C/D) are indicated. The nomenclature of the snoRNA was derived by homology to *L. major* snoRNA as described in (Eliaz *et al.,* RNA Bio. 2015). snoRNA levels verified by Northern analysis are highlighted in green and in yellow snoRNAs guiding hypermodified Ψs in LP parasites. The nomenclature identifies the snoRNAs via it chromosomal location and cluster location and its type (i.e. LD11Cs1H1 is H/ACA located on chromosome 11, the first cluster and the first snoRNA in the cluster).

| Chr. | Type | snoRNA | Length | EP rep1 | EP rep2 | LP rep1 | LP rep2 | Rep1 FC | Rep2 FC |
| --- | --- | --- | --- | --- | --- | --- | --- | --- | --- |
| 11 | HACA | LD11Cs1H1 | 69 | 43 | 41 | 396 | 184 | 9.21 | 4.49 |
| 11 | HACA | LD11Cs2H1 | 78 | 20 | 40 | 18 | 13 | 0.90 | 0.33 |
| 14 | CD | LD14Cs1C1 | 107 | 1183 | 546 | 752 | 320 | 0.64 | 0.59 |
| 14 | CD | LD14Cs1C2 | 112 | 2516 | 2520 | 6224 | 4457 | 2.47 | 1.77 |
| 14 | CD | LD14Cs1C3 | 80 | 1591 | 889 | 1485 | 647 | 0.93 | 0.73 |
| 14 | HACA | LD14Cs1H1 | 94 | 1034 | 562 | 2157 | 1546 | 2.09 | 2.75 |
| 14 | HACA | LD14Cs1H2 | 166 | 3048 | 2140 | 1950 | 1228 | 0.64 | 0.57 |
| 14 | HACA | LD14Cs1H3 | 283 | 4439 | 3188 | 1876 | 1359 | 0.42 | 0.43 |
| 15 | CD | LD15Cs1C1 | 104 | 164 | 371 | 209 | 171 | 1.27 | 0.46 |
| 15 | HACA | LD15Cs1H1 | 63 | 27 | 70 | 33 | 29 | 1.22 | 0.41 |
| 16 | HACA | LD16Cs1H1 | 68 | 8 | 3 | 6 | 1 | 0.75 | 0.33 |
| 16 | HACA | LD16Cs2H1 | 60 | 24 | 34 | 20 | 16 | 0.83 | 0.47 |
| 18 | CD | LD18Cs1C1 | 84 | 4433 | 3161 | 12870 | 6601 | 2.90 | 2.09 |
| 18 | CD | LD18Cs1C2 | 92 | 6366 | 5705 | 16116 | 11088 | 2.53 | 1.94 |
| 18 | CD | LD18Cs1C3 | 97 | 8427 | 7739 | 20350 | 13261 | 2.41 | 1.71 |
| 18 | HACA | LD18Cs1H1 | 68 | 893 | 613 | 4282 | 2270 | 4.80 | 3.70 |
| 18 | HACA | LD18Cs1H2 | 70 | 1155 | 460 | 2530 | 1141 | 2.19 | 2.48 |
| 19 | HACA | LD19Cs1H1 | 59 | 20 | 47 | 20 | 28 | 1.00 | 0.60 |
| 19 | HACA | LD19Cs1H2 | 76 | 24 | 29 | 18 | 18 | 0.75 | 0.62 |
| 19 | HACA | LD19Cs2H1 | 59 | 13 | 20 | 11 | 15 | 0.85 | 0.75 |
| 20 | CD | LD20Cs1C1 | 113 | 3696 | 3288 | 8345 | 5104 | 2.26 | 1.55 |
| 20 | CD | LD20Cs1C2 | 125 | 7436 | 5260 | 16047 | 12207 | 2.16 | 2.32 |
| 20 | CD | LD20Cs1C3 | 89 | 2587 | 2035 | 7154 | 3216 | 2.77 | 1.58 |
| 20 | CD | LD20Cs1C4 | 90 | 9029 | 7588 | 20810 | 12617 | 2.30 | 1.66 |
| 20 | CD | LD20Cs1C5 | 78 | 8374 | 6383 | 25047 | 17997 | 2.99 | 2.82 |
| 20 | HACA | LD20Cs-1H1 | 79 | 13 | 19 | 9 | 6 | 0.69 | 0.32 |
| 20 | CD | LD20Cs2C1 | 117 | 4143 | 2460 | 9635 | 5591 | 2.33 | 2.27 |
| 22 | CD | LD22Cs1C1 | 102 | 104 | 144 | 57 | 30 | 0.55 | 0.21 |
| 22 | CD | LD22Cs1C2 | 99 | 7910 | 8351 | 29350 | 23367 | 3.71 | 2.80 |
| 22 | HACA | LD22Cs-1H1 | 84 | 18 | 44 | 23 | 13 | 1.28 | 0.30 |
| 23 | CD | LD23Cs1C1 | 90 | 2226 | 1567 | 13956 | 10696 | 6.27 | 6.83 |
| 23 | CD | LD23Cs1C2 | 83 | 7530 | 7014 | 45608 | 30539 | 6.06 | 4.35 |
| 23 | CD | LD23Cs1C3 | 93 | 10066 | 9070 | 30803 | 25717 | 3.06 | 2.84 |
| 23 | HACA | LD23Cs1H1 | 65 | 901 | 497 | 4589 | 1767 | 5.09 | 3.56 |
| 23 | HACA | LD23Cs1H2 | 61 | 26844 | 12688 | 53895 | 21376 | 2.01 | 1.68 |
| 25 | CD | LD25Cs1C1 | 91 | 4022 | 3545 | 11481 | 8856 | 2.85 | 2.50 |
| 25 | CD | LD25Cs-1C1 | 102 | 60 | 123 | 83 | 129 | 1.38 | 1.05 |
| 25 | CD | LD25Cs1C2 | 82 | 4243 | 4113 | 10361 | 11776 | 2.44 | 2.86 |
| 25 | CD | LD25Cs1C3 | 76 | 23000 | 11412 | 31648 | 15984 | 1.38 | 1.40 |
| 25 | CD | LD25Cs1C4 | 114 | 23522 | 16073 | 50241 | 32958 | 2.14 | 2.05 |
| 26 | CD | LD26Cs1C1 | 99 | 11664 | 7058 | 31384 | 20288 | 2.69 | 2.87 |
| 26 | CD | LD26Cs1C2 | 75 | 710 | 648 | 3073 | 1451 | 4.33 | 2.24 |
| 26 | CD | LD26Cs1C3 | 119 | 6286 | 9595 | 20625 | 12164 | 3.28 | 1.27 |
| 26 | HACA | LD26Cs1H1 | 69 | 933 | 624 | 5603 | 2447 | 6.01 | 3.92 |
| 26 | HACA | LD26Cs1H2 | 81 | 3579 | 2108 | 8203 | 5834 | 2.29 | 2.77 |
| 26 | HACA | LD26Cs1H3 | 82 | 3570 | 2279 | 5802 | 2914 | 1.63 | 1.28 |
| 26 | HACA | LD26Cs1H4 | 68 | 11664 | 5943 | 26451 | 13335 | 2.27 | 2.24 |
| 26 | HACA | LD26Cs1H5 | 81 | 4868 | 3092 | 22273 | 16824 | 4.58 | 5.44 |
| 26 | HACA | LD26Cs1H6 | 70 | 2862 | 1389 | 3814 | 2713 | 1.33 | 1.95 |
| 26 | HACA | LD26Cs1H7 | 74 | 11353 | 5404 | 24950 | 15329 | 2.20 | 2.84 |
| 26 | HACA | LD26Cs1H8 | 66 | 402 | 294 | 2053 | 845 | 5.11 | 2.87 |
| 26 | HACA | LD26Cs1H9 | 69 | 2289 | 1061 | 2886 | 2199 | 1.26 | 2.07 |
| 26 | HACA | LD26Cs1H-9 | 87 | 1798 | 916 | 2548 | 1264 | 1.42 | 1.38 |
| 26 | CD | LD26Cs2C1 | 103 | 2874 | 2233 | 10013 | 7069 | 3.48 | 3.17 |
| 26 | HACA | LD26Cs2H1 | 91 | 18 | 31 | 25 | 14 | 1.39 | 0.45 |
| 26 | HACA | LD26Cs2H2 | 86 | 4151 | 2476 | 15920 | 11371 | 3.84 | 4.59 |
| 27 | CD | LD27Cs1C1 | 89 | 4177 | 2573 | 10352 | 13185 | 2.48 | 5.12 |
| 27 | CD | LD27Cs1C2 | 86 | 899 | 745 | 3493 | 7147 | 3.89 | 9.59 |
| 27 | CD | LD27Cs1C3 | 89 | 12290 | 9398 | 48768 | 33615 | 3.97 | 3.58 |
| 27 | HACA | LD27Cs1H1 | 66 | 1282 | 1060 | 11755 | 7880 | 9.17 | 7.43 |
| 27 | HACA | LD27Cs1H-1 | 62 | 671 | 561 | 1049 | 1266 | 1.56 | 2.26 |
| 27 | HACA | LD27Cs1H2 | 69 | 639 | 575 | 1000 | 1202 | 1.56 | 2.09 |
| 27 | HACA | LD27Cs1H3 | 69 | 4142 | 2318 | 11928 | 8104 | 2.88 | 3.50 |
| 27 | HACA | LD27Cs1H4 | 69 | 245 | 247 | 1671 | 1117 | 6.82 | 4.52 |
| 29 | HACA | LD29Cs1H1 | 181 | 2509 | 2945 | 6078 | 3990 | 2.42 | 1.35 |
| 29 | HACA | LD29Cs1H2 | 276 | 1207 | 912 | 866 | 640 | 0.72 | 0.70 |
| 29 | HACA | LD29Cs1H3 | 78 | 420 | 296 | 1554 | 862 | 3.70 | 2.91 |
| 29 | CD | LD29Cs2C1a | 101 | 1224 | 1082 | 2773 | 2321 | 2.27 | 2.15 |
| 29 | CD | LD29Cs2C1b | 101 | 1772 | 1377 | 5064 | 4715 | 2.86 | 3.42 |
| 30 | CD | LD30Cs1C1 | 89 | 4304 | 3357 | 12883 | 9494 | 2.99 | 2.83 |
| 30 | CD | LD30Cs1C2 | 102 | 6400 | 4797 | 16106 | 9703 | 2.52 | 2.02 |
| 30 | HACA | LD30Cs1'H1 | 75 | 132 | 51 | 338 | 169 | 2.56 | 3.31 |
| 30 | HACA | LD30Cs1'H2A | 108 | 2675 | 2148 | 6980 | 4917 | 2.61 | 2.29 |
| 30 | HACA | LD30Cs1'H2B | 103 | 6820 | 3861 | 10565 | 7515 | 1.55 | 1.95 |
| 30 | HACA | LD30Cs1'H3 | 65 | 1213 | 517 | 2520 | 1125 | 2.08 | 2.18 |
| 30 | CD | LD30Cs2C1 | 90 | 1901 | 1631 | 4379 | 4128 | 2.30 | 2.53 |
| 30 | CD | LD30Cs2C2 | 91 | 1879 | 1613 | 4330 | 4082 | 2.30 | 2.53 |
| 30 | HACA | LD30Cs2H1 | 75 | 698 | 344 | 2091 | 1457 | 3.00 | 4.24 |
| 30 | HACA | LD30Cs2H-1 | 73 | 502 | 238 | 1463 | 1241 | 2.91 | 5.21 |
| 30 | HACA | LD30Cs3H1 | 73 | 530 | 476 | 2210 | 1834 | 4.17 | 3.85 |
| 31 | CD | LD31Cs1C1 | 90 | 23653 | 18226 | 42189 | 46261 | 1.78 | 2.54 |
| 32 | CD | LD32Cs1C1 | 83 | 403 | 249 | 1032 | 909 | 2.56 | 3.65 |
| 32 | HACA | LD32Cs2H1 | 88 | 222 | 167 | 996 | 930 | 4.49 | 5.57 |
| 32 | HACA | LD32Cs3H1 | 220 | 2278 | 1758 | 2306 | 2183 | 1.01 | 1.24 |
| 33 | CD | LD33Cs1C1 | 95 | 22227 | 15052 | 62833 | 56785 | 2.83 | 3.77 |
| 33 | CD | LD33Cs1C2 | 92 | 5185 | 3038 | 11332 | 8610 | 2.19 | 2.83 |
| 33 | CD | LD33Cs1C3 | 103 | 37852 | 20779 | 63123 | 27628 | 1.67 | 1.33 |
| 33 | CD | LD33Cs1C4 | 102 | 47040 | 31838 | 141560 | 71567 | 3.01 | 2.25 |
| 33 | HACA | LD33Cs1H1 | 68 | 6259 | 3574 | 12704 | 6367 | 2.03 | 1.78 |
| 33 | HACA | LD33Cs1pH1 | 57 | 1652 | 2764 | 1230 | 1187 | 0.74 | 0.43 |
| 33 | HACA | LD33Cs1pH2 | 78 | 2016 | 2049 | 1707 | 1551 | 0.85 | 0.76 |
| 33 | CD | LD33Cs2C1 | 137 | 41929 | 16728 | 13786 | 5928 | 0.33 | 0.35 |
| 33 | CD | LD33Cs2C2 | 79 | 2080 | 1259 | 11250 | 5843 | 5.41 | 4.64 |
| 33 | HACA | LD33Cs2H1 | 70 | 3314 | 1809 | 8307 | 4680 | 2.51 | 2.59 |
| 33 | CD | LD33Cs3C1 | 113 | 13645 | 11437 | 30690 | 27950 | 2.25 | 2.44 |
| 33 | HACA | LD33Cs3H1 | 70 | 2717 | 1843 | 8330 | 4879 | 3.07 | 2.65 |
| 33 | HACA | LD33Cs3H2 | 60 | 2077 | 1364 | 12012 | 8445 | 5.78 | 6.19 |
| 33 | HACA | LD33Cs4H1 | 87 | 100 | 139 | 93 | 66 | 0.93 | 0.47 |
| 34 | CD | LD34Cs1C1 | 94 | 5639 | 5112 | 14720 | 15365 | 2.61 | 3.01 |
| 34 | CD | LD34Cs1C2 | 101 | 566 | 496 | 1451 | 1360 | 2.56 | 2.74 |
| 34 | HACA | LD34Cs1H1 | 75 | 538 | 327 | 1652 | 1074 | 3.07 | 3.28 |
| 34 | HACA | LD34Cs1H-1 | 73 | 83 | 41 | 284 | 141 | 3.42 | 3.44 |
| 34 | HACA | LD34Cs1H2 | 71 | 6806 | 4786 | 18598 | 11550 | 2.73 | 2.41 |
| 34 | HACA | LD34Cs2H2 | 69 | 88 | 44 | 300 | 149 | 3.41 | 3.39 |
| 34 | HACA | LD34Cs2H3 | 57 | 5 | 9 | 13 | 5 | 2.60 | 0.56 |
| 35 | CD | LD35Cs1C1 | 86 | 278 | 307 | 1571 | 1077 | 5.65 | 3.51 |
| 35 | CD | LD35Cs1C2 | 155 | 11702 | 5435 | 17010 | 8110 | 1.45 | 1.49 |
| 35 | HACA | LD35Cs1H1 | 79 | 435 | 320 | 2120 | 1463 | 4.87 | 4.57 |
| 35 | CD | LD35Cs2C1 | 79 | 4576 | 4214 | 7096 | 3716 | 1.55 | 0.88 |
| 35 | CD | LD35Cs2C2 | 85 | 3100 | 2281 | 5558 | 3076 | 1.79 | 1.35 |
| 35 | CD | LD35Cs2C3 | 140 | 5 | 6 | 5 | 3 | 1.00 | 0.50 |
| 35 | HACA | LD35Cs2pH1 | 80 | 18 | 32 | 19 | 17 | 1.06 | 0.53 |
| 35 | HACA | LD35Cs2pH2 | 78 | 19 | 24 | 33 | 16 | 1.74 | 0.67 |
| 35 | CD | LD35Cs3C1 | 105 | 2025 | 1293 | 4901 | 2344 | 2.42 | 1.81 |
| 35 | CD | LD35Cs3C-1 | 91 | 3102 | 1165 | 5713 | 3916 | 1.84 | 3.36 |
| 35 | CD | LD35Cs3C2 | 91 | 11782 | 8621 | 27107 | 31142 | 2.30 | 3.61 |
| 35 | CD | LD35Cs3C3 | 82 | 6138 | 5640 | 15394 | 13245 | 2.51 | 2.35 |
| 35 | CD | LD35Cs3C4 | 95 | 5056 | 3840 | 14380 | 10365 | 2.84 | 2.70 |
| 35 | CD | LD35Cs3C5 | 73 | 91086 | 71357 | 200136 | 150612 | 2.20 | 2.11 |
| 35 | CD | LD35Cs3C6 | 133 | 112287 | 56794 | 157294 | 91120 | 1.40 | 1.60 |
| 35 | CD | LD35Cs3C7 | 78 | 14606 | 13052 | 36506 | 24693 | 2.50 | 1.89 |
| 35 | HACA | LD35Cs3H1 | 73 | 2260 | 960 | 4431 | 1632 | 1.96 | 1.70 |
| 35 | HACA | LD35Cs3H2 | 73 | 2051 | 1149 | 6391 | 5654 | 3.12 | 4.92 |
| 35 | HACA | LD35Cs3H3 | 75 | 2447 | 1454 | 5862 | 3151 | 2.40 | 2.17 |
| 36 | CD | LD36Cs1C1 | 98 | 13227 | 7694 | 29803 | 18535 | 2.25 | 2.41 |
| 36 | CD | LD36Cs-1C1 | 98 | 1906 | 726 | 3845 | 3285 | 2.02 | 4.52 |
| 36 | CD | LD36Cs1C2 | 92 | 13610 | 10370 | 48701 | 22032 | 3.58 | 2.12 |
| 36 | CD | LD36Cs-1C2 | 86 | 5939 | 4587 | 19281 | 16676 | 3.25 | 3.64 |
| 36 | CD | LD36Cs1C3 | 108 | 7414 | 7156 | 23846 | 16520 | 3.22 | 2.31 |
| 36 | CD | LD36Cs1C4 | 130 | 2701 | 1842 | 8075 | 5747 | 2.99 | 3.12 |
| 36 | HACA | LD36Cs1H1 | 75 | 13113 | 8130 | 40134 | 31762 | 3.06 | 3.91 |
| 36 | HACA | LD36Cs-1H1 | 73 | 548 | 253 | 1451 | 1048 | 2.65 | 4.14 |
| 36 | HACA | LD36Cs1H2 | 72 | 3824 | 1825 | 5384 | 3286 | 1.41 | 1.80 |
| 36 | HACA | LD36Cs-1H2 | 68 | 33 | 36 | 34 | 14 | 1.03 | 0.39 |
| 36 | HACA | LD36Cs1H3 | 70 | 2229 | 1416 | 9074 | 3903 | 4.07 | 2.76 |
| 36 | HACA | LD36Cs1H4 | 65 | 4891 | 2509 | 7797 | 5124 | 1.59 | 2.04 |
| 36 | CD | LD36Cs1pC1 | 114 | 25 | 26 | 25 | 15 | 1.00 | 0.58 |
| 36 | CD | LD36Cs-1pC1 | 89 | 7571 | 3769 | 12715 | 8905 | 1.68 | 2.36 |
| 36 | CD | LD36Cs-1pC2 | 93 | 3878 | 3233 | 13528 | 10982 | 3.49 | 3.40 |
| 36 | HACA | LD36Cs1pH1 | 83 | 61 | 86 | 132 | 86 | 2.16 | 1.00 |
| 36 | HACA | LD36Cs-1pH1 | 65 | 2843 | 2254 | 17650 | 9251 | 6.21 | 4.10 |
| 36 | CD | LD36Cs2C1 | 106 | 35362 | 20623 | 51736 | 36851 | 1.46 | 1.79 |
| 36 | CD | LD36Cs2C-1 | 85 | 851 | 700 | 5396 | 2654 | 6.34 | 3.79 |
| 36 | CD | LD36Cs2C2 | 97 | 2414 | 2431 | 7578 | 4204 | 3.14 | 1.73 |
| 36 | CD | LD36Cs2C3 | 119 | 1660 | 1289 | 3718 | 3210 | 2.24 | 2.49 |
| 36 | HACA | LD36Cs2H1 | 72 | 9112 | 5286 | 24935 | 10883 | 2.74 | 2.06 |
| 36 | HACA | LD36Cs2pH1 | 191 | 6923 | 2443 | 7078 | 2941 | 1.02 | 1.20 |
| 36 | HACA | LD36Cs2ppH1 | 270 | 1646 | 1255 | 1033 | 758 | 0.63 | 0.60 |
| 36 | CD | LD36Cs3C1 | 90 | 7383 | 5165 | 15996 | 7989 | 2.17 | 1.55 |
| 36 | HACA | LD36Cs3H1 | 70 | 3815 | 2050 | 8460 | 8219 | 2.22 | 4.01 |
| 36 | HACA | LD36Cs3H-1 | 87 | 6006 | 3804 | 11751 | 6903 | 1.96 | 1.81 |
| 36 | HACA | LD36Cs3H2 | 57 | 3641 | 2574 | 6439 | 3406 | 1.77 | 1.32 |
| 36 | HACA | LD36Cs3H3 | 60 | 4257 | 3697 | 7872 | 4078 | 1.85 | 1.10 |
| 36 | HACA | LD36Cs3H4 | 76 | 1474 | 1085 | 4190 | 1822 | 2.84 | 1.68 |
| 36 | CD | LD36Cs4C1a | 146 | 2460 | 1895 | 3869 | 1776 | 1.57 | 0.94 |
| 36 | CD | LD36Cs4C1b | 138 | 1049 | 649 | 1658 | 689 | 1.58 | 1.06 |
| 36 | CD | LD36Cs4C2 | 135 | 91 | 147 | 87 | 58 | 0.96 | 0.39 |
| 36 | HACA | LD36Cs4H1a | 67 | 882 | 446 | 3227 | 2690 | 3.66 | 6.03 |
| 36 | HACA | LD36Cs4H1b | 70 | 844 | 427 | 3089 | 2574 | 3.66 | 6.03 |
| 36 | HACA | LD36Cs4H2 | 80 | 708 | 394 | 1458 | 1092 | 2.06 | 2.77 |
| 36 | HACA | LD36Cs5H1 | 80 | 90 | 54 | 197 | 137 | 2.19 | 2.54 |
| 4 | CD | LD4Cs1C1 | 154 | 3119 | 3661 | 12223 | 13032 | 3.92 | 3.56 |
| 4 | HACA | LD4Cs1H1 | 113 | 4029 | 2683 | 7564 | 5255 | 1.88 | 1.96 |
| 4 | HACA | LD4Cs1H2 | 76 | 908 | 410 | 2006 | 632 | 2.21 | 1.54 |
| 5 | CD | LD5Cs1C2 | 76 | 4699 | 2667 | 15397 | 18611 | 3.28 | 6.98 |
| 5 | CD | LD5Cs1C3 | 96 | 16287 | 10043 | 41506 | 34976 | 2.55 | 3.48 |
| 5 | CD | LD5Cs1C4 | 93 | 11417 | 6668 | 18456 | 17009 | 1.62 | 2.55 |
| 5 | CD | LD5Cs1C5 | 88 | 14796 | 8379 | 43543 | 48310 | 2.94 | 5.77 |
| 5 | HACA | LD5Cs1H1 | 289 | 3393 | 2398 | 2893 | 2727 | 0.85 | 1.14 |
| 5 | HACA | LD5Cs1H2 | 69 | 3228 | 2206 | 22938 | 12939 | 7.11 | 5.87 |
| 5 | HACA | LD5Cs1H3 | 75 | 497 | 262 | 1030 | 507 | 2.07 | 1.94 |
| 5 | HACA | LD5Cs1H4 | 69 | 2526 | 1188 | 4742 | 2777 | 1.88 | 2.34 |
| 8 | HACA | LD8Cs1H1 | 65 | 156 | 85 | 256 | 164 | 1.64 | 1.93 |
| 9 | HACA | LD9Cs1H1 | 65 | 50 | 67 | 48 | 18 | 0.96 | 0.27 |
