## Supplementary Data S1 for "Experimental evolution reveals post-transcriptional regulation as a novel driver of *Leishmania* fitness gain": combineGO.pl

```
#!/usr/bin/env perl
use strict;
use warnings;
use Pod::Usage;
use Getopt::Long;
use List::Util qw(first max maxstr min minstr reduce shuffle sum);
use Data::Dumper;
my $help = 0;
#print Dumper(\%go_ldbpk);
my @terms = ("CC","BP","MF");
#my @terms = ("BP");


#ld1s ge ids
my @ld1sGenes; open(F,"<Ld1S.ge.gtf") or die "cannot open Ld1S.ge.gtf\n$!\n"; while (<F>){  if($_=~/gene_id \"([^\"]+)\"/){ push(@ld1sGenes,$1); }} close F;

#initialize best go (storing the best GO for each gene) and best go info (storing the source of the best go identifier, for debugging)
my (%bestGo) ;
foreach my $term (@terms){
  foreach my $ld1s_ge (@ld1sGenes){
    $bestGo{$term}{$ld1s_ge}{"GOID"} = "NA";
    $bestGo{$term}{$ld1s_ge}{"info"} = "NA";
  }
}

#read GOID tables
my %go_ldbpk = loadGOIdDir("GO/tritrypGOtables/split_LdBPKv1","tritryp");
my %go_linf  = loadGOIdDir("GO/tritrypGOtables/split_Linfantum","tritryp");
my %go_lmaj  = loadGOIdDir("GO/tritrypGOtables/split_Lmajor","tritryp");
my %go_lmex  = loadGOIdDir("GO/tritrypGOtables/split_Lmexicana","tritryp");
my %go_tbru  = loadGOIdDir("GO/tritrypGOtables/split_Tbruceibrucei927","tritryp");
my %go_tcru  = loadGOIdDir("GO/tritrypGOtables/split_Tcruzi","tritryp");
my %go_ldbpk_uni = loadGOIdDir("uniprot","uniprot");


#melt ld1s ids (each Ld1s ge will point to the corresponding block of other identifiers, e.g. ldbpk, linf, tbru)
my %ld1sId2ldbpkBlock = loadEquivalenceTable("Ld1S_LdBPK_ID.equivalence.tsv");
my %ld1sId2linfBlock  = loadEquivalenceTable("GO/outOrthoFinder/Linfantum");
my %ld1sId2lmajBlock  = loadEquivalenceTable("GO/outOrthoFinder/Lmajor");
my %ld1sId2lmexBlock  = loadEquivalenceTable("GO/outOrthoFinder/Lmexicana");
my %ld1sId2tbruBlock  = loadEquivalenceTable("GO/outOrthoFinder/Tbruceibrucei927");
my %ld1sId2tcruBlock  = loadEquivalenceTable("GO/outOrthoFinder/Tcruzi");

#find GO
#NOTE1: if one Ld1S gene points to multiple target species genes (otherIdBlock), findGOID() will gather the non-redundant list of available GOs across all target genes
#NOTE2: findGOID() should be run sequentially in a hierarchical fashion, so that before you have the lower quality species and curation (e.g. Tcru computed) then you have increasingly higher confidence/quality GOIDs (e.g. Ldbpk curated) because each time you run it, if some GOIDs are found, they will replace those previously found.
findGOID(\%ld1sId2tcruBlock  , \%go_tcru  , "computed" , "Tcru");
findGOID(\%ld1sId2tbruBlock  , \%go_tbru  , "computed" , "Tbru");
findGOID(\%ld1sId2lmexBlock  , \%go_lmex  , "computed" , "Lmex");
findGOID(\%ld1sId2lmajBlock  , \%go_lmaj  , "computed" , "Lmaj");
findGOID(\%ld1sId2linfBlock  , \%go_linf  , "computed" , "Linf");
findGOID(\%ld1sId2ldbpkBlock , \%go_ldbpk , "computed" , "Ldbpk");
findGOID(\%ld1sId2tcruBlock  , \%go_tcru  , "curated" , "Tcru");
findGOID(\%ld1sId2tbruBlock  , \%go_tbru  , "curated" , "Tbru");
findGOID(\%ld1sId2lmexBlock  , \%go_lmex  , "curated" , "Lmex");
findGOID(\%ld1sId2lmajBlock  , \%go_lmaj  , "curated" , "Lmaj");
findGOID(\%ld1sId2linfBlock  , \%go_linf  , "curated" , "Linf");
findGOID(\%ld1sId2ldbpkBlock , \%go_ldbpk_uni , "uniprot" , "Ldbpk");
findGOID(\%ld1sId2ldbpkBlock , \%go_ldbpk , "curated" , "Ldbpk");

#manually assign GOIDs to ncRNAs
my %nc;
open(F,"<Ld1S.ge.gtf") or die "cannot open Ld1S.ge.gtf\n$!\n"; 
while (<F>){  
  if($_=~/.*ncRNAhomology.*gene_id \"([^\"]+)\".*type \"snoRNA\"/){ 
    $nc{$1}{'BP'}="GO:0006396";
    $nc{$1}{'CC'}="GO:0005730";
  } elsif ($_=~/.*ncRNAhomology.*gene_id \"([^\"]+)\".*type \"UsnRNA\"/) {
    $nc{$1}{'BP'}="GO:0009301;GO:0008380";
  } elsif ($_=~/.*ncRNAhomology.*gene_id \"([^\"]+)\".*type \"SLRNA\"/) {
    $nc{$1}{'BP'}="GO:0045291";
  } elsif ($_=~/.*ncRNAhomology.*gene_id \"([^\"]+)\".*type \"7SL\"/) {
    $nc{$1}{'BP'}="GO:0006616";
    $nc{$1}{'CC'}="GO:0048500";
  }
} 
close F;

#fix the bestGO with manual nc info
foreach my $term (@terms){
  foreach my $ld1s_ge (@ld1sGenes){
    if (($bestGo{$term}{$ld1s_ge}{"GOID"} eq 'NA') && (defined $nc{$ld1s_ge}{$term})){
       $bestGo{$term}{$ld1s_ge}{"GOID"} = $nc{$ld1s_ge}{$term};
       $bestGo{$term}{$ld1s_ge}{"info"} = 'manual_curation_ncRNA';
    }
  }
}

#print best go, best go info and cytoscape
foreach my $term (@terms){
  open(F1,">GO/${term}_bestGO");
  open(F2,">GO/${term}_bestGOinfo");
  system("rm -rf GO/${term}_bestGO.cytoscape");
  open(F3,">>GO/${term}_bestGO.cytoscape");
  my $headerType;
  if ($term eq 'BP'){$headerType='Biological Process';}elsif($term eq 'MF'){$headerType='Molecular Function';}elsif($term eq 'CC'){$headerType='Cellular Component';}
  print F3 "(species=Leishmania donovani)(type=${headerType})(curator=GO)\n";
  foreach my $ld1s_ge (@ld1sGenes){
    print F1 $ld1s_ge . "\t" . $bestGo{$term}{$ld1s_ge}{"GOID"} . "\n";
    print F2 $ld1s_ge . "\t" . $bestGo{$term}{$ld1s_ge}{"info"} . "\n";
    if ($bestGo{$term}{$ld1s_ge}{"GOID"} ne 'NA'){
      my @fi = split(/;/ , $bestGo{$term}{$ld1s_ge}{"GOID"});
      foreach my $go (@fi){
        $go=~s/GO://;
        print F3 "$ld1s_ge = $go\n";
      }
    }
  }
  close F1;
  close F2;
  close F3;
}


###########
#functions#
###########
sub loadGOIdDir {
  my ($functionDir , $source) = @_;
  my %h;
  foreach my $term (@terms){
    if ($source eq "tritryp"){
      open(CURATED ,"<${functionDir}/${term}_curated") or die "cannot open ${functionDir}/${term}_curated\n$!\n";
      open(COMPUTED,"<${functionDir}/${term}_computed") or die "cannot open${functionDir}/${term}_computed\n$!\n";
      while (<CURATED>){ if($_=~/(\S+)\s+(.+)/){ $h{$term}{"curated"}{$1} = $2;}} close CURATED;
      while (<COMPUTED>){ if($_=~/(\S+)\s+(.+)/){ $h{$term}{"computed"}{$1} = $2;}} close COMPUTED;
    } elsif ($source eq "uniprot"){
      open(UNI ,"<${functionDir}/${term}_uniprot") or die "cannot open ${functionDir}/${term}_uniprot\n$!\n";
      while (<UNI>){ if($_=~/(\S+)\s+(.+)/){ $h{$term}{"uniprot"}{$1} = $2;}} close UNI;
    }
  }
  return %h;
}

sub loadEquivalenceTable {
  my ($equivalenceTable) = @_;
  my %h;
  open(F,"<$equivalenceTable") or die "cannot open $equivalenceTable\n$!\n";
  my $dumpHeader = <F>;
  while (<F>){
    if($_=~/(\S+)\s+(\S+)/){
      my $ld1sBlock    = $1;
      my $otherIdBlock = $2;
      next if ($ld1sBlock eq "NA");
      my @ld1sBlockArray    = split(/,/, $ld1sBlock);
      my @otherIdBlockArray = split(/,/, $otherIdBlock);
      foreach my $ld1s_ge (@ld1sBlockArray){
        @{$h{$ld1s_ge}} = @otherIdBlockArray;
      }
    }
  }
  close F;
  return %h;
}

sub findGOID {
  my %ld1sId2otherIdBlock = %{$_[0]};
  my %go_hash  = %{$_[1]};
  my $curation = $_[2];
  my $info     = $_[3];

  foreach my $term (@terms){
    foreach my $ld1s_ge (@ld1sGenes){
      my %allGoPerGe;
      foreach my $other_ge (@{$ld1sId2otherIdBlock{$ld1s_ge}}){
        if ($other_ge ne "NA") {
         if ($go_hash{$term}{$curation}{$other_ge}){
           my $go = $go_hash{$term}{$curation}{$other_ge} ;
           $allGoPerGe{$go} = 1;
         }
        }
      }
      my $gos =join(';',keys(%allGoPerGe));
      $gos=~s/(;)?N\/A//g;
      $gos=~s/^;//;
      $gos=~s/^;//;
      if($gos ne ""){
        $bestGo{$term}{$ld1s_ge}{"GOID"} = $gos ;
        $bestGo{$term}{$ld1s_ge}{"info"} = $info . "_" . $curation ;
      }
    } 
  }
}
```
