## Supplementary Data S1 for "Experimental evolution reveals post-transcriptional regulation as a novel driver of *Leishmania* fitness gain": run.sh

```
#!/bin/bash
#Get GO descriptions for BP, MF andd CL terms (so 3 files listing genes and GO IDs)
#GO will follow a hierarchical attribution:
#GO tritrypdb curated > GO GAF > GO homology curated > GO tritryp inferred > GO homology inferred 

#get proteins (downloaded the 11092019, release 2019-09)
mkdir -p GO/proteins
cd GO/proteins
wget ftp://ftp.sanger.ac.uk/pub/genedb/releases/latest/Lmajor/Lmajor_proteins.fasta.gz
wget ftp://ftp.sanger.ac.uk/pub/genedb/releases/latest/Lmexicana/Lmexicana_proteins.fasta.gz
wget ftp://ftp.sanger.ac.uk/pub/genedb/releases/latest/Linfantum/Linfantum_proteins.fasta.gz
wget ftp://ftp.sanger.ac.uk/pub/genedb/releases/latest/Tbruceibrucei927/Tbruceibrucei927_proteins.fasta.gz
wget ftp://ftp.sanger.ac.uk/pub/genedb/releases/latest/Tcruzi/Tcruzi_proteins.fasta.gz
for X in `ls *proteins.fasta.gz`; do 
  N=`echo $X | sed -e 's/_proteins.fasta.gz/.pep.fa/'`
  zcat  $X | perl -ne 'if($_=~/>(.+):pep /){print ">$1\n"}else{print;}' > $N 
  rm -rf $X  
done
cd ..

#orthofinder
module load fasta mafft
module load fastme/2.1.6.1 
module load mcl/12-068
orthofinder=/pasteur/entites/HubBioIT/gio/apps/OrthoFinder-2.2.7/orthofinder
CPU=1
mkdir -p outOrthoFinder
for X in Linfantum Lmajor Lmexicana Tbruceibrucei927 Tcruzi ; do
  mkdir -p orthofinder_$X
  cp proteins/${X}.pep.fa orthofinder_$X
  zcat ../Ld1S.pep.fa.gz | perl -ne 'if($_=~/>.+ gene:(.+)/){print ">$1\n"}else{print;}' > orthofinder_$X/Ld1S.pep.fa
  $orthofinder -f orthofinder_$X -S diamond -t $CPU
  out=`find orthofinder_$X | grep Orthogroups.csv`
  cp $out outOrthoFinder/$X
  rm -rf orthofinder_$X
  cat outOrthoFinder/$X | cut -f 1  --complement > outOrthoFinder/_tmp
  mv outOrthoFinder/_tmp  outOrthoFinder/$X
done 

#get tritryp GO tables (downloaded the 11092019)
mkdir -p tritrypGOtables
for X in Linfantum Lmajor Lmexicana Tbruceibrucei927 Tcruzi ; do
  grep ">" proteins/${X}.pep.fa | tr -d ">" > tritrypGOtables/${X}.ids
done
cut -f 4 /pasteur/projets/policy01/BioIT/Giovanni/datasets/projects/p2p5/LdBPKv1.ge.bed > tritrypGOtables/LdBPKv1.ids
#THEN I GO HERE: http://tritrypdb.org/tritrypdb/showQuestion.do?questionFullName=GeneQuestions.GeneByLocusTag  and I MANUALLY download all annotations
mv GeneByLocusTag_Summary.txt Linfantum.tsv
mv GeneByLocusTag_Summary.txt Lmajor.tsv
mv GeneByLocusTag_Summary.txt Lmexicana.tsv
mv GeneByLocusTag_Summary.txt Tbruceibrucei927.tsv
mv GeneByLocusTag_Summary.txt Tcruzi.tsv
mv GeneByLocusTag_Summary.txt LdBPKv1.tsv
#remove spaces from header
cd /pasteur/projets/policy01/BioIT/Giovanni/datasets/assemblies/leishmanias/peterMyler/GO/tritrypGOtables
sed -i '1s/ /_/g' *.tsv
cd ..
#split tsv
Rscript ../split_tritrypGOtables.R
#find best GO_ID for each gene
cd ..
perl combineGO.pl
```
