## Supplementary Data S1 for "Experimental evolution reveals post-transcriptional regulation as a novel driver of *Leishmania* fitness gain": split_tritrypGOtables.R

```
allSpecies <- gsub( x=list.files("/pasteur/projets/policy01/BioIT/Giovanni/datasets/assemblies/leishmanias/peterMyler/GO/tritrypGOtables/",pattern=".tsv") , pattern=".tsv" , replacement="")
for (species in allSpecies) {
  outDir <- paste0("/pasteur/projets/policy01/BioIT/Giovanni/datasets/assemblies/leishmanias/peterMyler/GO/tritrypGOtables/split_",species)
  system (paste0("mkdir -p ", outDir))
  df <- read.delim(paste0("/pasteur/projets/policy01/BioIT/Giovanni/datasets/assemblies/leishmanias/peterMyler/GO/tritrypGOtables/",species,".tsv"))
  write.table(df[,c("Input_ID","Computed_GO_Component_IDs")] ,quote=F , sep="\t" , row.names=F , col.names=F , file=paste0(outDir,"/CC_computed"))
  write.table(df[,c("Input_ID","Computed_GO_Function_IDs")]  ,quote=F , sep="\t" , row.names=F , col.names=F , file=paste0(outDir,"/MF_computed"))
  write.table(df[,c("Input_ID","Computed_GO_Process_IDs")]   ,quote=F , sep="\t" , row.names=F , col.names=F , file=paste0(outDir,"/BP_computed"))
  write.table(df[,c("Input_ID","Curated_GO_Component_IDs")]  ,quote=F , sep="\t" , row.names=F , col.names=F , file=paste0(outDir,"/CC_curated"))
  write.table(df[,c("Input_ID","Curated_GO_Function_IDs")]   ,quote=F , sep="\t" , row.names=F , col.names=F , file=paste0(outDir,"/MF_curated"))
  write.table(df[,c("Input_ID","Curated_GO_Process_IDs")]    ,quote=F , sep="\t" , row.names=F , col.names=F , file=paste0(outDir,"/BP_curated"))
}
```
